# Identification of T-Cell specific signature miRNAs and target pathways in Psoriasis

**DOI:** 10.1101/2025.02.16.638502

**Authors:** Priyanka Madaan, Nipanshi Tyagi, Sourabh Tyagi, Hemant Ritturaj Kushwaha, Ravi Tandon, Diya A’gitok Sangma, Manju Jain

## Abstract

Psoriasis is a chronic skin disease characterized by keratinocyte-hyperproliferation, involving T cells as the major players in the manifestation of lesional and systemic inflammation. With the extensive exploration of keratinocyte-dysfunctionality, T cell-centric deregulation has gained much focus with limited molecular understanding in the context of disease etiology. miRNAs, the molecular-switches known to drive disease manifestation via altering keratinocyte biology, have not been much studied in the context of T cell alterations. This study aims to explore T cell-associated miRNAs and their target signaling networks to understand their role in modulating psoriasis-specific T cell changes. miRNAs with altered expression in psoriatic T cells were identified through a comprehensive analysis of the available literature and datasets. The relevant miRNAs were used for prediction and enrichment of cognate-targets and mapping of associated pathways involved in different modalities of T cell functionality. The study identified 14 miRNAs dysregulated in psoriatic T cells along with 256 downstream targets involved in 15 distinct T cell-associated signaling pathways. 5/14 miRNAs identified were further validated with a significant increase in their expression in enriched CD3^+^T cells from blood of psoriatic patients versus healthy subjects. The psoriatic T cell specific miRNA-driven signaling cascades identified were modelled to propose yet unexplored molecular basis of dysregulated T cell activation, proliferation, survival, and subset differentiation associated with the disease. The new findings delineate candidate miRNA–target signaling components that need to be validated in T cell disease etiology. Importantly, the key miRNAs/targets can be employed towards psoriasis-specific diagnostics and therapeutic interventions.

## 1. Background

Psoriasis is an autoimmune skin disease characterized with typical red, itchy, scaly skin lesions [1]. Its worldwide occurrence is 2-3% [2]. The disease exhibits multiple phenotypes according to distribution, anatomical localization, size and nature of lesions, disease onset, and severity. Clinically designated psoriasis types include Plaque-type/Psoriasis vulgaris (PV), guttate, erythrodermis and generalized pustular psoriasis with PV as the most prevalent form [3]. It is an outcome of deregulated crosstalk between keratinocytes and immune cells [4]. The skin cells/Keratinocytes exhibit hyperproliferation, parakeratosis, and acanthosis, giving rise to typical psoriatic lesions with infiltration of immune cells. Alteration in keratinocyte biology has been explored much to understand the disease etiology, albeit the role of immune cells with T cells as major players in disease progression and activity has taken center stage more recently [5]. Distinct T cell subsets viz. Th1, Th17, and Th22, with their effector cytokines, have been reported to play a key role in establishing an inflammatory landscape, as evident from the successful use of biologics viz. etanercept, infliximab, secukinumab in disease treatment [4, 6]. Deciphering molecular mechanisms of T cell dysregulation is an important area that needs to be explored further to unravel T cell-specific immune checkpoints involved in disease etiology and can provide potential diagnostic markers and therapeutic targets. microRNAs (miRNAs), comprising small non-coding RNA molecules involved in the modulation of cellular gene expression, represent critical molecular regulators in the pathogenesis of psoriasis [7, 8]. A large number of differentially expressed miRNAs have been reported in psoriasis patients compared to healthy individuals in various sample types viz skin lesions, PBMCs, T cells, blood, plasma, and serum [9–13]. Based on the differential expression profile in patients, specific miRNAs have been reported as potential diagnostic and prognostic biomarkers in relation to disease severity and clinical efficacy of treatments [14–17]. A significant body of literature has demonstrated miRNA-driven downstream mechanisms leading to dysregulation of keratinocyte proliferation and differentiation culminating in altered skin cycle and development of psoriatic lesions [13, 18–20]. Unlike keratinocyte-associated miRNAs, fewer reports are available on T cell-specific miRNAs, limited to reporting their altered expression pattern with only four miRNAs worked out for their downstream target axes [10, 21–26].

This study puts together a comprehensive meta-analysis of miRNAs with altered expression in psoriatic T cells. Using *in silico* approaches, the candidate T cell-associated miRNAs retrieved have been worked out for their downstream targetome with an attempt to decipher the potential T cell signaling pathways affected in disease condition. The molecular signaling network potentially modulable via differential expression of select miRNAs in psoriatic T cells has been further modeled to propose the molecular basis of psoriasis-associated alterations in T cell activation, proliferation, survival and differentiation. In line with the huge knowledge gap on miRNA- mediated T cell dysregulation mechanisms in psoriasis, our findings provide a set of key T cell miRNAs and their target axes that should be pursued for their functional role in disease biology with the potential to develop T-cell specific diagnostic biomarkers and therapeutic strategies.

## 2. Methods

### 2.1 Selection of miRNAs associated with T cells in psoriasis

A literature search was performed for “microRNAs Psoriasis T cells” and “miRNAs Psoriasis T cells” using PubMed, Web of Science, and Scopus databases. The same search terms were used to extract GEO datasets related to miRNAs in T cells in psoriasis (Table **S1**). Based on the literature and datasets available, relevant miRNAs with altered expression in psoriatic T cells were selected for further work, as shown in **Fig. 1**.

**Fig. 1.**
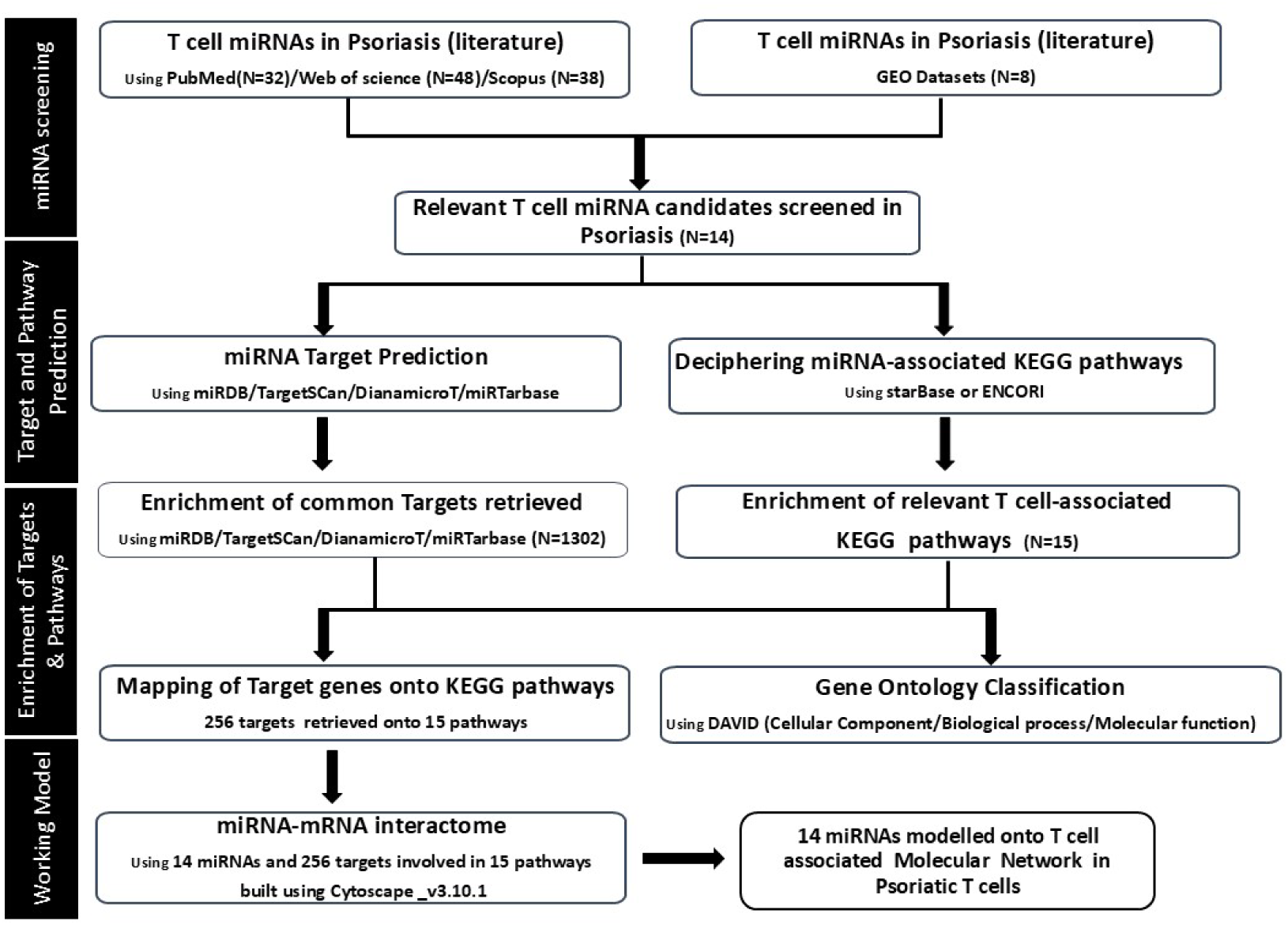
Flow Chart with detailed Methodology used for target prediction and pathway analysis for 14 miRNAs relevant in Psoriatic T cells.

#### Expression analysis of select miRNAs

Blood samples from clinically confirmed psoriasis vulgaris patients and healthy control subjects were used to enrich CD3+ T cells. Briefly, ficoll density gradient was used to retrieve PBMCs, that were further processed to isolate CD3+ T cells employing positive selection based Magnetic separation with Human CD3 MicroBeads (130-050-101) and Miletnyi MiniMACS starting Kit (Cat.no 130-090-312) as per the manufacturers’ instructions. miRNA isolation from enriched T cells (psoriasis, n=12; control, n=12) was performed using miRNeasy Mini Kit (Qiagen, Cat.no 217004) as per standard protocol. 300ng of RNA was used to synthesized cDNA using miRCURY LNA RT Kit (Qiagen, Cat. No 339340) according to the manufacturer’s instructions. cDNA was used for qRT-PCR using MiRCURY LNA SYBR Green PCR Kit (Qiagen Cat. No. 339345) and primers for 5 select miRNAs namely: miR-223: 339306 (YP00205986); miR-142-3p: 339306 (YP00204291); miR-155: MS00031486; miR-146a: MS00003535; miR-21: MS00009086. The study received formal ethical approval from AIIMS Bathinda (IEC/AIIMS/BTI/150), and the Central University of Punjab (CUPB/IEC/2016/045) with written informed consent obtained from all subjects.

### 2.2 Prediction and enrichment of miRNA-specific targets and pathways

4 distinct miRNA target prediction tools were employed to predict target genes for 14 miRNAs of interest: miRDB (http://www.mirdb.org/<u>), TargetScan</u> (https://www.targetscan.org/vert_80/), DianamicroT (https://bio.tools/DIANA-microT) and mirTarBase (https://mirtarbase.cuhk.edu.cn/). Venny 2.0 was utilized to identify common targets across the four databases for each of the miRNAs. Simultaneously, KEGG (Kyoto Encyclopedia of Genes and Genomes) pathways associated with the 14 miRNAs of interest were retrieved using starBase or ENCORI (https://starbase.sysu.edu.cn/). Relevant T cell associated KEGG pathways were filtered from the total complement of the KEGG pathways enriched. The common targets downstream of 14 miRNAs retrieved from the 4 miR-target prediction tools were then overlapped with components of selected T cell- associated KEGG pathways. Targets involved in the resultant T cell-associated pathways were mapped for their interaction network using Cytoscape _v3.10.1.

### 2.3 Gene ontology (GO) analysis of targets

GO enrichment analysis was accomplished using David (Database for Annotation, Visualization, and Integrated Discovery), an integrated biological knowledge base and analytic tool aimed at systematically extracting the biological significance of gene sets (https://david.ncifcrf.gov/; version 6.8). The target gene list was given as input, with biological, cellular, and molecular annotations as the output data.

### 2.4 Generation of Working Model

The key T cell-associated pathways involving relevant miRNA-target axes deciphered for psoriasis were mapped for the overlaps. The resulting miR-target correlates were assessed in line with T cell modalities known to be dysregulated in psoriasis as per literature viz altered T cell activation, proliferation, survival and altered T cell subsets exhibiting enhanced Th1/Th17 subset differentiation. The model projects the potential miRNA- mediated molecular mechanisms that need to be explored to understand T-cell disease etiology and generate a pool of novel biomarkers and therapeutic targets.

## 3. Results

### 3.1 Psoriasis specific screening of differentially expressed T cell-associated miRNAs

A comprehensive literature search was performed, employing PUBMED, Web of Science, and Scopus for the search strings “microRNAs Psoriasis T cells” and “miRNAs psoriasis T cells.” Along with the literature search, the Gene Expression Omnibus (GEO) database was also searched for these strings to retrieve relevant datasets. As depicted in **Fig. 1**, a total of 32 articles were retrieved from PubMed, 48 articles were retrieved from Web of Science, and 38 articles from Scopus. After removing the overlapping literature, a total of 38 publications, comprising 12 review articles and 26 research studies, were retrieved. A thorough assessment of reviews with their cited references and the research articles gave a total of 8 relevant research studies. 7 out of the 8 studies focused on specific miRNAs with their downstream effector mechanism in T cells/T cell subsets with only a single article with miRNA profiling of sorted T cell subsets from psoriasis patients [10, 21–27]. With respect to the GEO database searched, a total of 8 datasets were retrieved **(Table S1)**. Only 1 out of 8 datasets related to T cell-associated miRNA expression pattern could be derived based on profiling of dermal infiltrates from psoriasis subjects with qRT-PCR validation of the candidate T cell/T cell subset specific miRNAs by Lovendorf et al. (2015) (GEO dataset: GSE 57012) [10]. The remaining datasets represented transcriptomic studies on lesional skin samples with no useful information on T cell-specific miRNAs. A total of 14 miRNAs with differential expression in T cells/T cell subsets alongside other sample types, viz. skin, PBMCs, serum, and plasma, were narrowed down with significance in psoriasis. miRNA-specific expression changes for the 14 miRNAs as per sample type with their reported/potential role in T cell-specific phenomena in psoriasis are detailed in **Table 1**. Expression pattern validation of 5/14 miRNAs was done using CD3+ T cells enriched from blood samples of psoriasis versus healthy subjects. All the 5 miRNAs *viz* miR-223, miR-142-3p, miR-155, miR146a and miR21 showed a significant upregulation in psoriatic T cells (**Fig 2**).

**Fig. 2.**
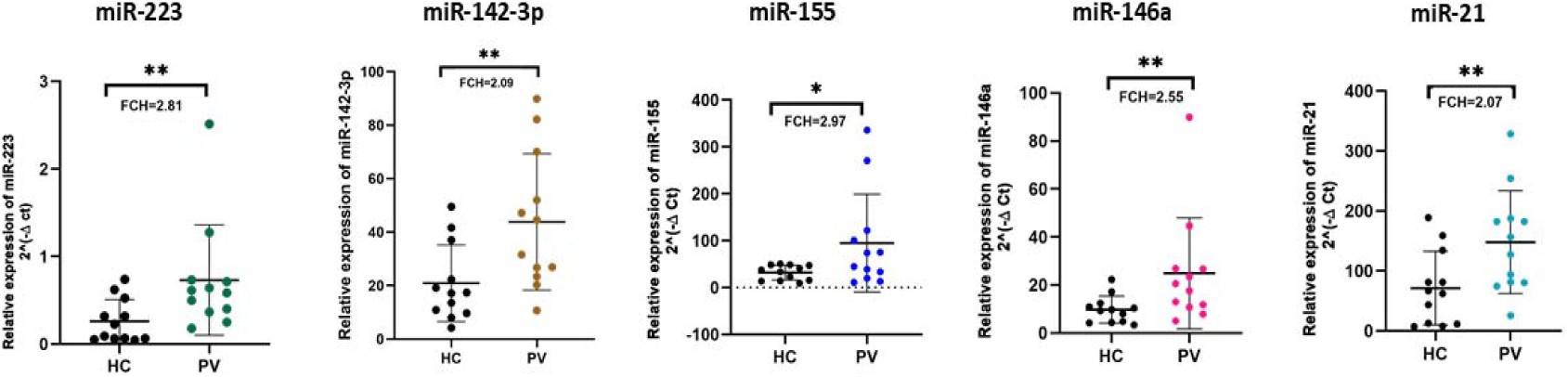
Expression analysis of 5 miRNAs: miR-223, miR-142-3p, miR-155, miR-146a and miR-21 exhibited upregulation in psoriatic T cells compared to healthy controls. miR-223 (Fold change = 2.81, p-value <0.01), miR-142-3p (Fold change = 2.09, p-value <0.01), miR-155 (Fold change = 2.97, p-value <0.05), miR-146a (Fold change =2.55, p-value <0.01) and miR-21 (Fold change = 2.07, p-value <0.05). Enriched CD3+ T cells with >95 % purity from psoriatic patients and healthy control group were used to check expression level of these 5 miRNAs.

**Table 1.** miRNAs relevant to T cells in Psoriasis.

| Sr. No | miRNAs | Sample type | Expression | Role in Psoriasis | Targets | References |
|---|---|---|---|---|---|---|
| 1. | miR-138<sup>#</sup> | CD4<sup>+</sup> T cells | Downregulated | Th1/ Th2 balance | RUNX3 | [21, 28] |
|  |  | Skin, HACAT cells | Downregulated | Cell proliferation | hTERT |  |
| 2. | miR-210<sup>#</sup> | CD4<sup>+</sup> T cells | Upregulated | Regulates Th subset (Th1/Th17/Tregs) Differentiation | STAT6, LYN and FoXP3 | [22-24, 29] |
|  |  | Skin lesions, KCs Dermal infiltrates, PBMC, Mice models, serum | Upregulated | Role in the development of skin lesions | Not reported |  |
| 3. | miR-200a<sup>#</sup> | CD4<sup>+</sup> T cells | Upregulated | Correlates with Th17/Treg cells and their cytokines | Not reported | [25] |
| 4. | let -7a<sup>#</sup> | T cells | Downregulated | T cell Proliferation | STAT3 | [26] |
|  |  | Skin lesion | Downregulated | NA |  |  |
| 5. | miR-193b* | Th1 and PBMCs | Upregulated | Potential role in maintaining T cell naivety | Not reported | [10, 11, 24] |
|  |  | Th17 cells | Downregulated | Potentially regulates Th17 Differentiation | Not reported |  |
| 6. | miR-21* | T cells | Upregulated | T cell survival | Not reported | [9, 10, 27, 30-32] |
|  |  | Th17 subsets | Downregulated | Potentially regulates Th17/Treg balance | Not reported |  |
|  |  | Skin lesions, KCs, Dermal infiltrates, PBMCs, Mice Model, Plasma, Serum | Upregulated | Regulates epidermal hyperplasia in psoriasis | TIMP3 (KCs) |  |
| 7. | miR-223* | Naïve T cells, Tregs | Downregulated | Potential role in T cell activation and Th17 differentiation | Not reported | [10, 11, 31, 33] |
|  |  | Th17 cells | Upregulated |  |  |  |
|  |  | Skin lesions, Dermal infiltrates PBMCs, Plasma | Upregulated | Promotes KC proliferation under IL-22 stimulation inhibits apoptosis | PTEN |  |
| 8. | miR-99a* | Naïve T cells, Th1 cells | Upregulated | High expression with T cells naïvity | Not reported | [10, 19, 34] |
|  |  | Tregs | Downregulated | NA |  |  |
|  |  | Skin lesion HACAT cell, Dermal infiltrates, PBMCs | Downregulated | Regulates Keratinocyte proliferation and differentiation | IGF1R, FZD5/FZD8 |  |
| 9. | miR-125b* | Naïve T cells | Upregulated | Potential role in maintaining T cell Naivety | Not Reported | [9, 10, 35-37] |
|  |  | Treg cells | Downregulated | Regulating Tregs |  |  |
|  |  | Skin lesions, KCs, dermal infiltrates, PBMCs | Downregulated | KC proliferation | BRD4, FGFR2, USP2 |  |
| 10. | miR 17-92*<br>(cluster) | T cells | Upregulated (miR-17, miR-19a)<br>Downregulated (miR-92) | Potentially regulates T cell subset differentiation | Not reported | [38, 39] |
|  |  | Skin lesion, KCs | Upregulated | Lesion development Proliferation of KCs, increase in chemokine production in KCs | CDKN2B, SOCS1 |  |
| 11. | miR-146a* | Th17 cells | Downregulated | Potential Role in T cell activation and differentiation | Not reported | [10, 31, 40, 41] |
|  |  | Skin lesions, Dermal infiltrates PBMCs, serum | Upregulated | Positively correlates with IL-17A through IRAK1 and maintains inflammatory Response | IRAK1 |  |
|  |  | Plasma samples | Varied reports (Up/Down) | NA |  |  |
| 12. | miR-155* | Naïve T cells/T cell subsets | No change (small sample size) | Potential role in T cell activation and Th subset differentiation | Not reported | [10, 29, 31, 42, 43] |
|  |  | Skin lesions, HACAT, Dermal infiltrates, Dermal MSCs, PBMCs, Plasma, Serum | Upregulated | Promotes keratinocyte growth and inflammatory responses | NLRP3 |  |
| 13. | miR-142-3p* | T cells | No change (small sample size) | Potential role in T cell activation and differentiation | Not reported | [10, 44-46] |
|  |  | Skin Lesion Dermal infiltrates | Upregulated | Regulates KC proliferation and apoptosis | Sema3A |  |
|  |  | Serum | Varying reports | NA |  |  |
| 14. | miR-150* | Skin lesions, KCs, Dermal infiltrates | Downregulated | Inhibits KC viability and proliferation in hypoxic Conditions | HIF-1α and VEGFA | [10, 47] |

\# Target-specific validation in T cells reported for 4/14 miRNAs.
\* For 10/14 miRNAs, only expression profiling has been checked in T cells/T cell subsets in psoriasis patients vs controls with no validation work with respect to the molecular targets affected. The miRNA-specific targets given in respective rows are specific to the sample type.

### 3.2. Elucidation of T cell-specific miRNA targets, pathways, and functions

#### 3.2.1. Target and pathway prediction

Target mRNAs for the 14 miRNAs of interest were retrieved using 4 Target prediction tools detailed in the materials and methods section. Overall, a total of 1302 targets as overlaps from 4 prediction tools were retrieved for the 14 miRNAs as depicted in the Venn diagrams (Supplementary File S1). Concurrently, downstream KEGG pathways for the 14 miRNAs were enriched using starBase or ENCORI. The irrelevant pathways, such as KC-associated pathways viz. melanogenesis, neurotrophin signaling pathways, actin cytoskeleton signaling pathway, and several disease-associated pathways viz. prostate cancer and small cell lung cancer pathways were excluded in further analysis. Relevant T cell-associated pathways like the TCR signaling pathway, Th1/Th2/Th17 differentiation, IL-17 signaling pathway, and PI3K-AKT/MAPK/mTOR pathway were focused with the identification of a total of 15 pathways enlisted in Table 2. The 15 enriched pathways were mapped for their components against all the 1302 miRNA targets, with the identification of 256/1302 meaningful targets (Table 2 and Supplementary Table S2). Table 2 displays the 15 pathways identified involving one or multiple given targets downstream of 14 miRNAs, while Table S2 reports miR 17-92 cluster-specific additional targets.

**Table 2.**
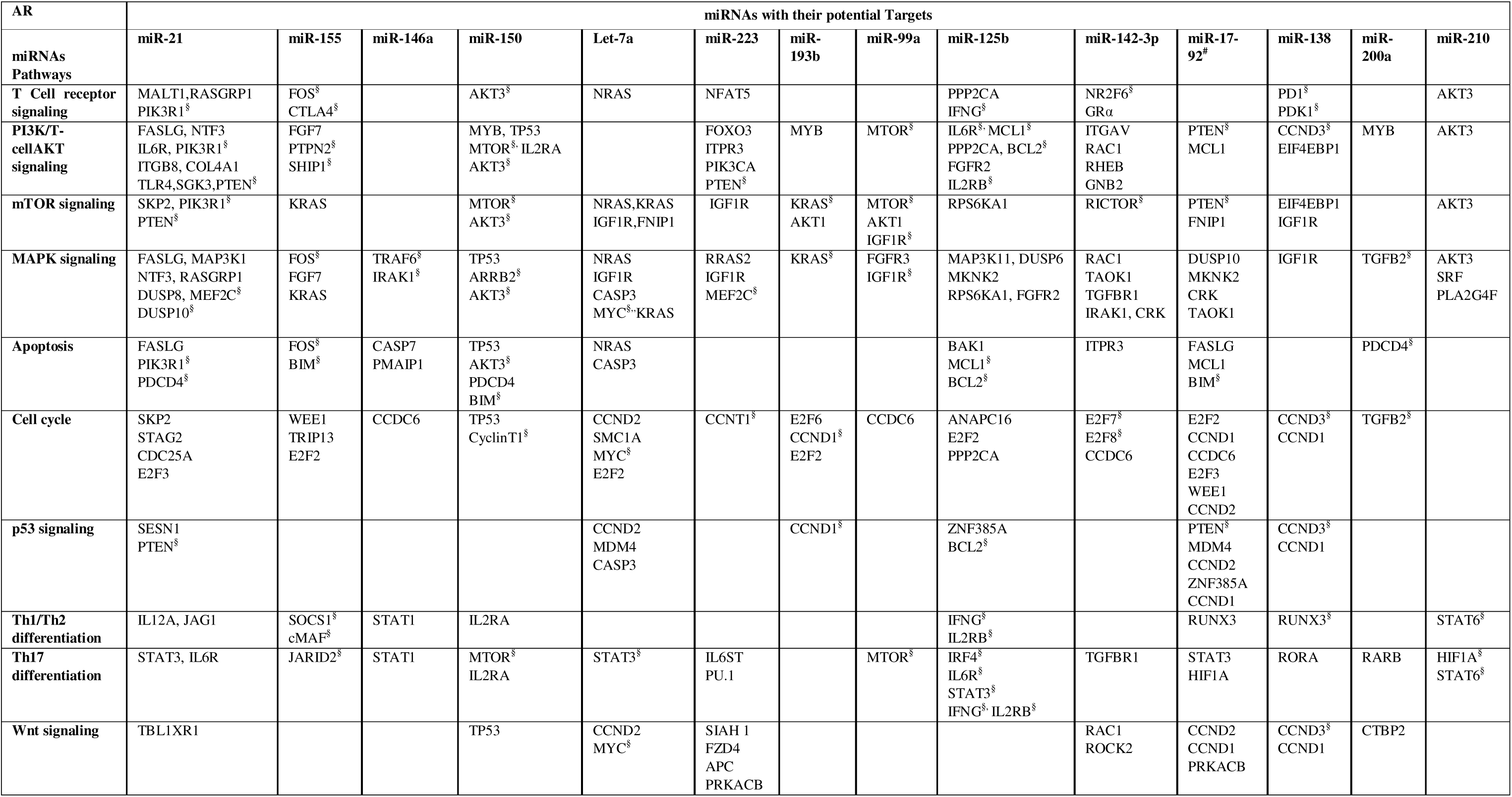
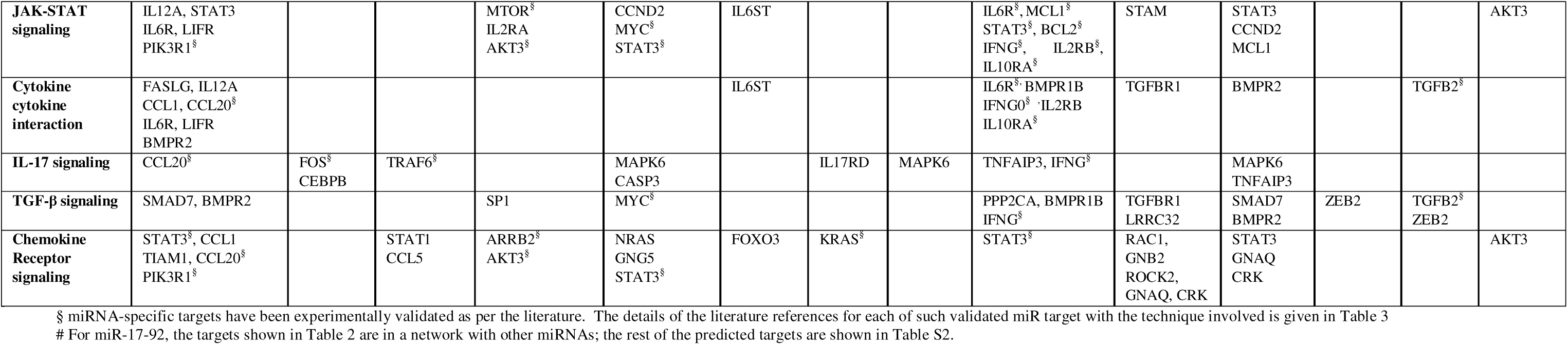
T cell-associated miRNAs in psoriasis with their potential downstream targets and the pathways deciphered using target prediction tools. miRNA targets superscribed with § have also been experimentally validated.

| **AR** | **miRNAs with their potential Targets** |  |  |  |  |  |  |  |  |  |  |  |  |  |
|---|---|---|---|---|---|---|---|---|---|---|---|---|---|---|
| **miRNAs<br>Pathways** | **miR-21** | **miR-155** | **miR-146a** | **miR-150** | **Let-7a** | **miR-223** | **miR-193b** | **miR-99a** | **miR-125b** | **miR-142-3p** | **miR-17-92<sup>#</sup>** | **miR-138** | **miR-200a** | **miR-210** |
| **T Cell receptor signaling** | MALT1,RASGRP1<br>PIK3R1<sup>§</sup> | FOS<sup>§</sup><br>CTLA4<sup>§</sup> |  | AKT3<sup>§</sup> | NRAS | NFAT5 |  |  | PPP2CA<br>IFNG<sup>§</sup> | NR2F6<sup>§</sup><br>GRα |  | PD1<sup>§</sup><br>PDK1<sup>§</sup> |  | AKT3 |
| **PI3K/T-cellAKT signaling** | FASLG, NTF3<br>IL6R, PIK3R1<sup>§</sup><br>ITGB8, COL4A1<br>TLR4,SGK3,PTEN<sup>§</sup> | FGF7<br>PTPN2<sup>§</sup><br>SHIP1<sup>§</sup> |  | MYB, TP53<br>MTOR<sup>§,</sup> IL2RA<br>AKT3<sup>§</sup> |  | FOXO3<br>ITPR3<br>PIK3CA<br>PTEN<sup>§</sup> | MYB | MTOR<sup>§</sup> | IL6R<sup>§,</sup> MCL1<sup>§</sup><br>PPP2CA, BCL2<sup>§</sup><br>FGFR2<br>IL2RB<sup>§</sup> | ITGAV<br>RAC1<br>RHEB<br>GNB2 | PTEN<sup>§</sup><br>MCL1 | CCND3<sup>§</sup><br>EIF4EBP1 | MYB | AKT3 |
| **mTOR signaling** | SKP2, PIK3R1<sup>§</sup><br>PTEN<sup>§</sup> | KRAS |  | MTOR<sup>§</sup><br>AKT3<sup>§</sup> | NRAS,KRAS<br>IGF1R,FNIP1 | IGF1R | KRAS<sup>§</sup><br>AKT1 | MTOR<sup>§</sup><br>AKT1<br>IGF1R<sup>§</sup> | RPS6KA1 | RICTOR<sup>§</sup> | PTEN<sup>§</sup><br>FNIP1 | EIF4EBP1<br>IGF1R |  | AKT3 |
| **MAPK signaling** | FASLG, MAP3K1<br>NTF3, RASGRP1<br>DUSP8, MEF2C<sup>§</sup><br>DUSP10<sup>§</sup> | FOS<sup>§</sup><br>FGF7<br>KRAS | TRAF6<sup>§</sup><br>IRAK1<sup>§</sup> | TP53<br>ARRB2<sup>§</sup><br>AKT3<sup>§</sup> | NRAS<br>IGF1R<br>CASP3<br>MYC<sup>§,</sup>KRAS | RRAS2<br>IGF1R<br>MEF2C<sup>§</sup> | KRAS<sup>§</sup> | FGFR3<br>IGF1R<sup>§</sup> | MAP3K11, DUSP6<br>MKNK2<br>RPS6KA1, FGFR2 | RAC1<br>TAOK1<br>TGFBR1<br>IRAK1, CRK | DUSP10<br>MKNK2<br>CRK<br>TAOK1 | IGF1R | TGFB2<sup>§</sup> | AKT3<br>SRF<br>PLA2G4F |
| **Apoptosis** | FASLG<br>PIK3R1<sup>§</sup><br>PDCD4<sup>§</sup> | FOS<sup>§</sup><br>BIM<sup>§</sup> | CASP7<br>PMAIP1 | TP53<br>AKT3<sup>§</sup><br>PDCD4<br>BIM<sup>§</sup> | NRAS<br>CASP3 |  |  |  | BAK1<br>MCL1<sup>§</sup><br>BCL2<sup>§</sup> | ITPR3 | FASLG<br>MCL1<br>BIM<sup>§</sup> |  | PDCD4<sup>§</sup> |  |
| **Cell cycle** | SKP2<br>STAG2<br>CDC25A<br>E2F3 | WEE1<br>TRIP13<br>E2F2 | CCDC6 | TP53<br>CyclinT1<sup>§</sup> | CCND2<br>SMC1A<br>MYC<sup>§</sup><br>E2F2 | CCNT1<sup>§</sup> | E2F6<br>CCND1<sup>§</sup><br>E2F2 | CCDC6 | ANAPC16<br>E2F2<br>PPP2CA | E2F7<sup>§</sup><br>E2F8<sup>§</sup><br>CCDC6 | E2F2<br>CCND1<br>CCDC6<br>E2F3<br>WEE1<br>CCND2 | CCND3<sup>§</sup><br>CCND1 | TGFB2<sup>§</sup> |  |
| **p53 signaling** | SESN1<br>PTEN<sup>§</sup> |  |  |  | CCND2<br>MDM4<br>CASP3 |  | CCND1<sup>§</sup> |  | ZNF385A<br>BCL2<sup>§</sup> |  | PTEN<sup>§</sup><br>MDM4<br>CCND2<br>ZNF385A<br>CCND1 | CCND3<sup>§</sup><br>CCND1 |  |  |
| **Th1/Th2 differentiation** | IL12A, JAG1 | SOCS1<sup>§</sup><br>cMAF<sup>§</sup> | STAT1 | IL2RA |  |  |  |  | IFNG<sup>§</sup><br>IL2RB<sup>§</sup> |  | RUNX3 | RUNX3<sup>§</sup> |  | STAT6<sup>§</sup> |
| **Th17 differentiation** | STAT3, IL6R | JARID2<sup>§</sup> | STAT1 | MTOR<sup>§</sup><br>IL2RA | STAT3<sup>§</sup> | IL6ST<br>PU.1 |  | MTOR<sup>§</sup> | IRF4<sup>§</sup><br>IL6R<sup>§</sup><br>STAT3<sup>§</sup><br>IFNG<sup>§,</sup> IL2RB<sup>§</sup> | TGFBR1 | STAT3<br>HIF1A | RORA | RARB | HIF1A<sup>§</sup><br>STAT6<sup>§</sup> |
| **Wnt signaling** | TBL1XR1 |  |  | TP53 | CCND2<br>MYC<sup>§</sup> | SIAH 1<br>FZD4<br>APC<br>PRKACB |  |  |  | RAC1<br>ROCK2 | CCND2<br>CCND1<br>PRKACB | CCND3<sup>§</sup><br>CCND1 | CTBP2 |  |
| **JAK-STAT signaling** | IL12A, STAT3<br>IL6R, LIFR<br>PIK3R1<sup>§</sup> |  |  | MTOR<sup>§</sup><br>IL2RA<br>AKT3<sup>§</sup> | CCND2<br>MYC<sup>§</sup><br>STAT3<sup>§</sup> | IL6ST |  |  | IL6R<sup>§</sup>, MCL1<sup>§</sup><br>STAT3<sup>§</sup>, BCL2<sup>§</sup><br>IFNG<sup>§</sup>, IL2RB<sup>§</sup>, IL10RA<sup>§</sup> | STAM | STAT3<br>CCND2<br>MCL1 |  |  | AKT3 |
| **Cytokine cytokine interaction** | FASLG, IL12A<br>CCL1, CCL20<sup>§</sup><br>IL6R, LIFR<br>BMPR2 |  |  |  |  | IL6ST |  |  | IL6R<sup>§,</sup> BMPR1B<br>IFNG0<sup>§</sup> <sup>,</sup>IL2RB<br>IL10RA<sup>§</sup> | TGFBR1 | BMPR2 |  | TGFB2<sup>§</sup> |  |
| **IL-17 signaling** | CCL20<sup>§</sup> | FOS<sup>§</sup><br>CEBPB | TRAF6<sup>§</sup> |  | MAPK6<br>CASP3 |  | IL17RD | MAPK6 | TNFAIP3, IFNG<sup>§</sup> |  | MAPK6<br>TNFAIP3 |  |  |  |
| **TGF-β signaling** | SMAD7, BMPR2 |  |  | SP1 | MYC<sup>§</sup> |  |  |  | PPP2CA, BMPR1B<br>IFNG<sup>§</sup> | TGFBR1<br>LRRC32 | SMAD7<br>BMPR2 | ZEB2 | TGFB2<sup>§</sup><br>ZEB2 |  |
| **Chemokine Receptor signaling** | STAT3<sup>§</sup>, CCL1<br>TIAM1, CCL20<sup>§</sup><br>PIK3R1<sup>§</sup> |  | STAT1<br>CCL5 | ARRB2<sup>§</sup><br>AKT3<sup>§</sup> | NRAS<br>GNG5<br>STAT3<sup>§</sup> | FOXO3 | KRAS<sup>§</sup> |  | STAT3<sup>§</sup> | RAC1,<br>GNB2<br>ROCK2,<br>GNAQ, CRK | STAT3<br>GNAQ<br>CRK |  |  | AKT3 |

§ miRNA-specific targets have been experimentally validated as per the literature. The details of the literature references for each of such validated miR target with the technique involved is given in Table 3
\# For miR-17-92, the targets shown in Table 2 are in a network with other miRNAs; the rest of the predicted targets are shown in Table S2.

#### 3.2.2. Functional ontology of targets

To access the overall significance of miRNAs in relation to the 256 relevant targets, Gene Ontology (GO) functional annotation was initially done using DAVID-based analysis. The principal biological processes (BP) shown to be affected downstream of the target genes were apoptosis, transcription/transcription regulation, cell cycle, and host-virus interaction (Fig. 3). With partial relevance of biological processes enriched, the targets were primarily localized to the nucleus, cytoplasm, and plasma membrane as per cellular components enriched. Molecular function terms (MF) showed a predominance of signal transduction activating and inhibiting proteins like kinases, serine-threonine protein kinase, protein phosphatase, activator, and repressor. For an in-depth understanding of T cell-specific dysregulation via signaling networks, select miRNA-mRNA axes were delineated based on the miRNA-mRNA interaction network.

**Fig. 3.**
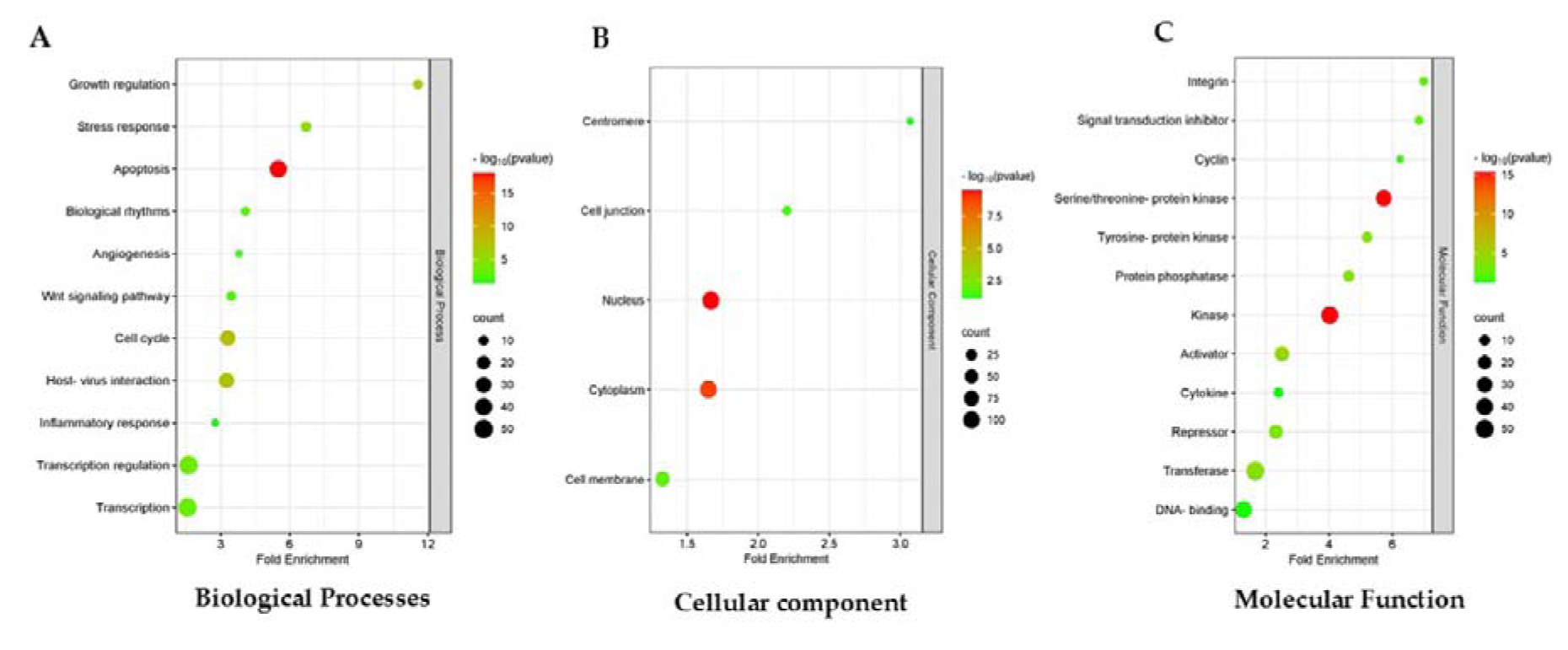
Functional Gene Ontology analysis was done for all the 256 targets downstream of 14 miRNAs with a role in T cells. DAVID functional GO analysis of (A) Eleven GO terms were deciphered in biological processes (B) Five GO terms enriched in cellular components (C) Twelve GO terms enriched in molecular function. The size of the circle indicates the number of genes, and color is associated with respective –log10^pvalue^.

### 3.3. Understanding miRNA-mRNA dysregulation in psoriatic T cells

The miRNA-mRNA interactome network was constructed using Cytoscape _v3.10.1 with miRNAs of interest as input shown in rectangular nodes and their mRNA targets represented as pathway-specific-colored spherical nodes (Fig. **4**). The T cell miRNAs with their targets exhibited an interconnected network representing 15 pathways relevant in the context of T cell functionality viz T cell activation, proliferation, apoptosis, and/or differentiation (Fig. 4 and Table 2). All the select miRNAs can target multiple pathways involving one or more than one target. Multiple miRNAs are seen to impinge on the same target as well, while a single miRNA holds the potential to regulate multiple targets with different signaling outcomes. Some of the miRNAs could affect specific signaling axes via unique targets. With this landscape of complex regulatory crosstalk, a set of targets downstream of the miR-17-92 cluster enlisted in Supplementary Table S2 are exclusive to six members of this cluster, unlike those shown in Table **2**, albeit affecting the same pathways. All the 15 pathways involving targets downstream of 14 miRNAs altered in psoriatic T cells can play a role in T cell etiology of disease. Most pathway-specific targets comprise predicted candidates, with only a few experimentally validated molecules in the context of different sample types and healthy/disease subjects (Table **2** and Table **3**). The miRNA- mRNA target axes and the composite pathway network altered in a psoriatic T cell are detailed in the discussion section.

**Fig. 4.**
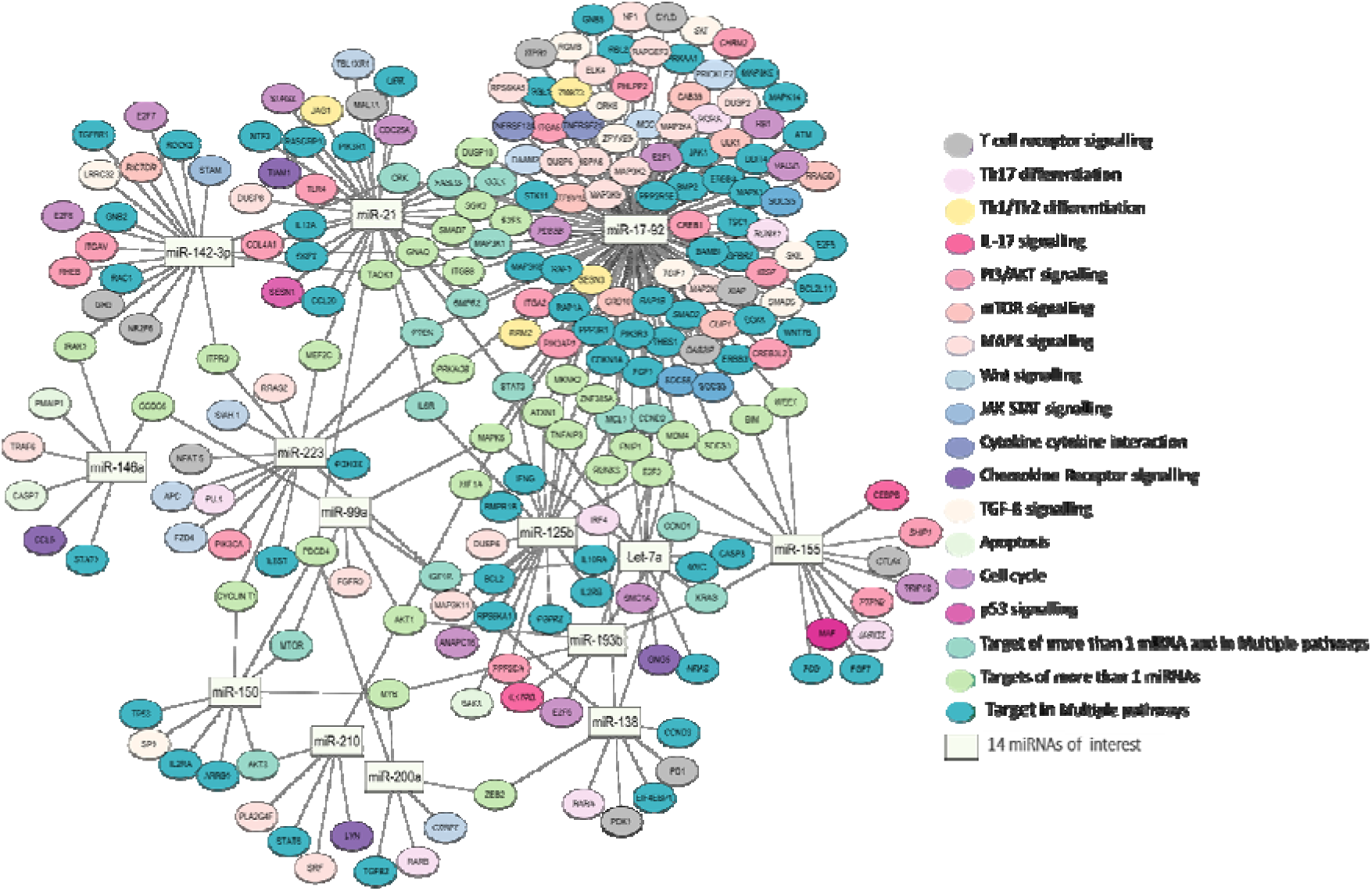
miRNA-mRNA interactome showing 14 miRNAs with their respective targets relevant in psoriatic T cells. The miRNA-specific target components are color-coded as per their involvement in 15 different signaling pathways potentially dysregulated in Psoriatic T cells. Rectangle nodes represent miRNAs, and spherical nodes represent mRNA targets.

**Table 3.** miRNAs with their literature-based validated targets.

| Sr. No | miRNA | Target | Sample Type/ disease | Techniques used in Target Validation (Approximate fold change values deciphered from published data*) |  |  | References |
|---|---|---|---|---|---|---|---|
|  |  |  |  | Reporter assay | Western blotting | qRT-PCR |  |
| 1. | miR-142-3p | RICTOR | CD4<sup>+</sup> T cells/ Aspergillus fumigatus infection |  | FCH≈ −2.5 | FCH≈ −2.5 | [48] |
|  |  | NR2F6 | NSCLC cell line<br>Lung adenocarcinoma | Binding checked with dual luciferase reporter assay | FCH≈ −2.0 | FCH≈ −2.0 | [49] |
|  |  | E2F7 | T cells (mice model) | Binding checked with Luciferase assay |  | FCH≈ −6.0 | [50] |
|  |  | E2F8 | T cells (mice model) | Binding checked with Luciferase assay |  | FCH≈ −6.0 | [50] |
| 2. | miR-150 | Cyclin T1 | CD4<sup>+</sup> T cells<br>Healthy subjects and Hela cell line |  |  | FCH≈ −10 | [51] |
|  |  | mTOR | CD4<sup>+</sup> T cells/<br>Mice model | Binding checked with Luciferase assay |  |  | [52] |
|  |  | AKT3/Bim | CD4<sup>+</sup> T cells<br>Healthy subjects | Binding checked with Luciferase assay |  |  | [53] |
|  |  | ARRB2 | CD4<sup>+</sup> T cells<br>Healthy subjects | Binding checked with Luciferase assay |  |  | [54] |
| 3. | miR-99a | IGF1R | HACAT cell line/<br>Psoriasis | Binding checked with Luciferase assay | FCH≈ −5.0 |  | [19] |
|  |  | mTOR | CD4<sup>+</sup> T cells/<br>Mice model | Binding checked with Luciferase assay |  |  | [52] |
| 4. | miR-223 | PTEN | HACAT cell line/<br>Psoriasis | Binding checked with dual luciferase assay | FCH≈ −2.5 | FCH≈ −3.3 | [33] |
|  |  | Cyclin T1 | CD4<sup>+</sup> T cells<br>Healthy subjects and Hela cell line |  |  | FCH≈ −10 | [51] |
| 5. | miR-138 | PD1 | CD4<sup>+</sup> T cells<br>Healthy subjects | Binding checked with luciferase assay |  |  | [55] |
|  |  | PDK1 | A549 cell line<br>NSCLC | Binding checked with Luciferase assay |  | FCH ≈ −3.5 | [56] |
|  |  | RUNX3 | CD4<sup>+</sup> T cells/<br>Psoriasis | Binding checked with Luciferase assay | FCH≈ −3.3 | FCH≈ −2.0 | [21, 57] |
|  |  | CCND3 | HCC tissue/ cell line<br>Hepatocellular carcinoma | Binding checked with dual luciferase reporter assay | FCH≈ −2.2 |  | [57] |
| 6. | miR-210 | LYN | CD4<sup>+</sup>T cells/Psoriasis | Binding checked with Luciferase assay | FCH≈ −3.8 |  | [23] |
|  |  | STAT-6 | CD4<sup>+</sup> T cells/Psoriasis | Binding checked with Luciferase assay | FCH≈ −25 |  | [23] |
|  |  | HIF1a | CD4<sup>+</sup> T cells/ Mice model,<br>hypoxic condition | Binding checked with Luciferase assay |  |  | [58] |
| 7. | Let-7a | STAT-3 | CD4<sup>+</sup> T cells/Psoriasis | Binding checked with Luciferase assay | FCH≈−0.3 |  | [26] |
|  |  | c-MYC | Namalwa cells/<br>Burkitt Lymphoma |  | FCH≈ −1.3 | FCH≈ 1.42 | [59] |
| 8. | miR-155 | SOCS1 | CD4<sup>+</sup> T cells<br>Mice model |  | FCH≈ −2.0 |  | [60] |
|  |  | c-Maf | Innate lymphoid cells /Mice Model Rhinitis | Binding checked with Luciferase assay | FCH≈ −2.5 | FCH≈ −1.5 | [61] |
|  |  | JARID2 | CD4<sup>+</sup> T cells<br>Mice model |  | FCH≈ −2.0 |  | [62] |
|  |  | CTLA4 | PBMCs<br>Atopic dermatitis | Binding checked with Luciferase assay |  |  | [63] |
|  |  | PTPN2<br>SHIP-1 | CD8<sup>+</sup> T cells/<br>Mice model | Binding checked with Luciferase assay | FCH≈ −2.0 |  | [64] |
|  |  | PIK3R1 | T cells/<br>Healthy subjects |  |  | FCH≈ −2.0 | [65] |
|  |  | cFOS | Mouse dendritic cell line | Binding checked with Luciferase assay |  | FCH≈ −2.5 | [66] |
|  |  | BIM | Jurkat cells/ CD4<sup>+</sup> T cells mice model |  | FCH≈ −2.5 |  | [67] |
| 9. | miR-21 | PIK3R1 | Human breast cancer cell lines | Binding checked with Luciferase assay | FCH≈ −3.0 | FCH≈ −3.0 | [68] |
|  |  | PDCD4 | Glioblastoma cell lines |  | FCH≈ −3.0 |  | [69] |
|  |  | CCL20 | Colorectal cancer cell lines | Binding checked with Luciferase assay | FCH≈ −2.5 | FCH≈ −2.5 | [70] |
|  |  | PTEN | Hepatocellular Carcinoma cells | Binding checked with Luciferase assay |  |  | [71] |
|  |  | MEF2C | Neuronal cells/<br>human SIV/HIV neurological disease | Binding checked with Luciferase assay |  |  | [72] |
|  |  | DUSP10 | CD8<sup>+</sup> T cells/<br>Healthy donors | Binding checked with Luciferase assay |  | FCH≈ −6.0 | [73] |
| 10. | miR-193b | KRAS | Human esophageal squamous cell Carcinoma | Binding checked with Luciferase assay |  |  | [74] |
|  |  | MYB | T-cell acute lymphoblastic leukemia | Binding checked with Luciferase assay |  |  | [75] |
|  |  | CCND1 | Prostate cancer cell lines | Binding checked with Luciferase assay | FCH ≈ −1.2 |  | [76] |
| 11. | miR-146a | TRAF6<br>IRAK1 | Macrophage cell lines | Binding checked with Luciferase assay |  |  | [77] |
| 12. | miR-200a | TGFB2 | Renal cell carcinoma | Binding checked with Luciferase assay |  |  | [78] |
|  |  | PCDCD4 | K562 cell line<br>Erythropoiesis differentiation study | Binding predicted, validated via knockout study |  |  | [79] |
| 13. | miR-17-92 | PTEN<br>BIM | Mice model<br>(Autoimmunity) | Binding checked with Luciferase assay |  |  | [80] |
| 14. | miR-125b-5p | IL6R<br>BCL-2<br>MCL-1 | Hepatocellular Carcinoma cell lines | Binding checked with Luciferase assay |  |  | [81] |
|  |  | IFNG<br>IL10RA<br>IL2RB | CD4<sup>+</sup> T cells<br>Healthy subjects | Binding checked with Luciferase assay |  |  | [82] |
|  |  | IRF4 | 293 T cell line<br>Macrophage study | Binding checked with Luciferase assay |  |  | [83] |
|  |  | STAT3 | CD3<sup>+</sup> T cells<br>Systemic lupus | Binding checked with Luciferase assay |  |  | [84] |

\*The techniques used for validation, along with the binding specificity and fold changes, as elucidated from the data published in cited references, are presented for each of the miRNAs of interest.

### 3.4 miRNAs as molecular switches dysregulating Psoriatic T cells

With knowledge of 14 miRNAs that are differentially expressed in psoriatic T cells along with their specific targets, working models have been put together warranting experimental exploration to understand the miRNA- mediated molecular mechanisms underlining altered T cell activation/proliferation/ survival and differentiation.

#### 3.4.1 T cell activation miR-target axes in psoriasis

Our findings on T cell-specific miRNAs with altered expression in psoriasis, along with their elucidated targets superimposed with key pathways involved in T cell activation/survival/proliferation, depict the potential miRNA-target axes that can dysregulate psoriatic T cells as shown in Fig. 5 Panel A and Panel B. Altered miRNAs depicted as rectangular boxes are shown to target the cognate validated molecules, shown in solid lines while the predicted molecules are shown to be targeted as dotted lines.

**Fig. 5.**
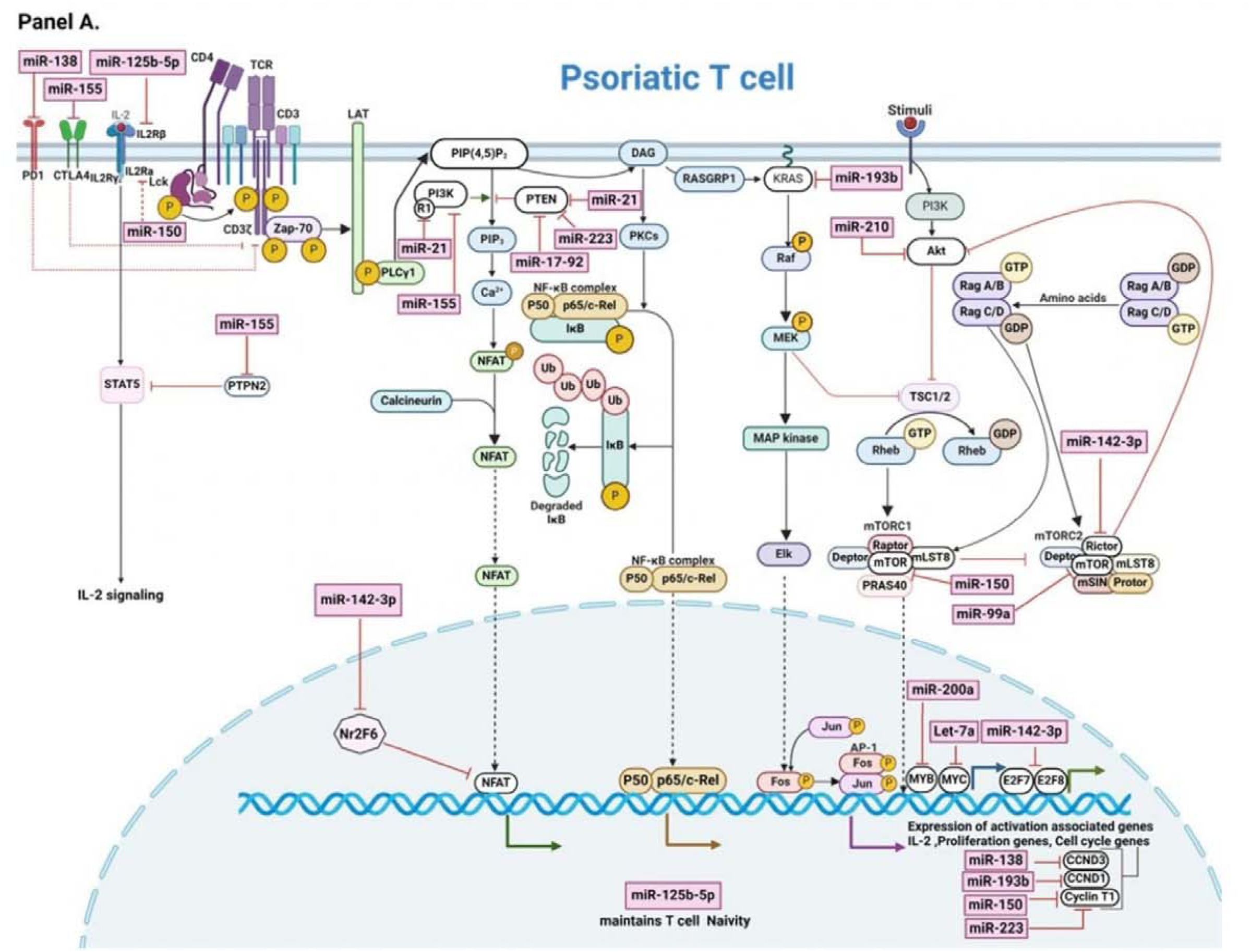
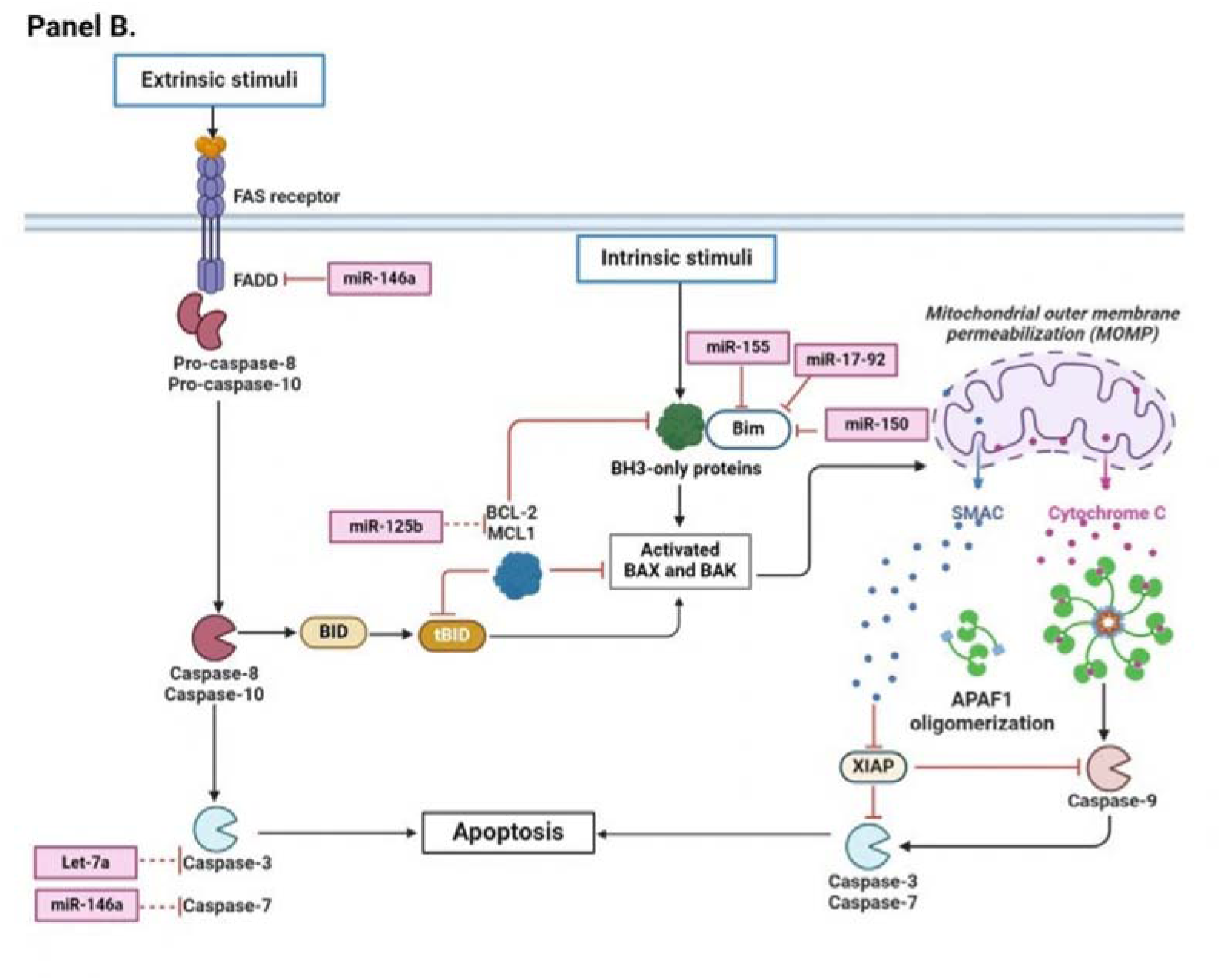
Panel. **A**. Working Model for miRNAs deciphered for their role in altered T cell activation, proliferation, and apoptosis. miRNAs are represented in rectangular boxes with solid lines representing validated targets and dotted lines representing predicted targets,14 miRNAs with a role in psoriatic T cells are shown to impinge downstream targets that make a part of T cell activation and proliferation pathways (TCR signaling, PI3K signaling, MAPK signaling, and mTOR signaling) (Figure created in BioRender. Madaan, P. (2024) https://BioRender.com/e43o810). **Panel B**. Pathways involved in apoptosis (extrinsic and intrinsic) involving 6 /14 miRNAs are shown alongside (Figure created in BioRender. Madaan, P. (2025) https://BioRender.com/j22v025)

The differential expression of components comprising key activation TCR axis with associated housekeeping signaling cascades PI3K/AKT, mTOR, MAPK, and NF-kB along with the major transcription factors viz. NFAT/ NFKB/ AP-1, interacting partners like MYB, MYC and E2Fs, and downstream readouts of IL-2, cell cycle, and proliferation-associated genes can mediate psoriasis-specific T cell changes. The miRNA-driven effector modalities, PI3K/AKT cascade can be altered by miR-21, miR-155, miR-223, and miR-17-92, mTOR pathway can be altered by miR-210, miR-142-3p, miR-150, and miR-99a and MAPK signaling can be dysregulated by miR-193b along with the inhibitory cascades CTLA4 and PD1 that can be modulated by miR- 155 and miR-138 respectively in psoriasis. T cell activation-associated transcription factors and their downstream effector genes represent another layer of molecules that can be dysregulated in psoriasis as miR- 193b, let-7a, and miR-142-3p can alter the expression of the transcription factors MYB, MYC, and E2Fs while miR-138, miR-193b, miR-150, and miR-223 can potentially alter the level of cyclins viz. CCND3, CCND1 and CyclinT1. miR-150 can also potentially alter the IL-2 signaling axis via IL2RA, a predicted target deciphered in our study. Modulation of pro-apoptotic molecules like BIM, Caspase 3, and Caspase 7, and anti-apoptotic molecules like BCL-2 and MCL-1 can also facilitate psoriatic T cell phenotype, as depicted in Fig. 5 Panel B. The miRNA-target coordinates mapped in Fig. 5 Panel A and Panel B provide valuable insights into the potential molecular axes that can modulate the activation and proliferation status of psoriatic T cells.

#### 3.4.2. T cell differentiation miR-target axes in psoriasis

Overlap of relevant miRNA-target axes retrieved with components of T cell differentiation pathways reveal potential molecular changes associated with Th1/Th2 balance and Th17 differentiation status in psoriasis as illustrated in Fig. 6 Panel A and Panel B. Multiple validated and few of the predicted targets of 9/14 miRNAs constitute components of Th1/Th2/Th17 subset specific differentiation pathways known to be altered in psoriasis. miR-138, miR-210, and Let-7a and the corresponding validated targets in psoriatic T cells, imply their significance in disease etiology. With these proof-of-concept miR-targets retrieved in our findings, miR-125b downregulated in differentiated T cells can potentially promote Th1 and Th17 differentiation in disease condition by mediating the enhanced expression of Th1-inducing IFN-γ, IL2RB, and Th17-specific IL6R, STAT-3 and IRF4. Similarly, miR-155 targeting Jarid2 and c-Maf, miR-146a-TRAF6, miR-210-HIF1α, and miR-200a-TGFβ2 as validated findings along with predicted miR-target checkpoints depicted as dotted lines suggest multiple miR-target axes that can modulate T cell differentiation towards a Th1/Th17 dominant response in psoriasis.

**Fig. 6.**
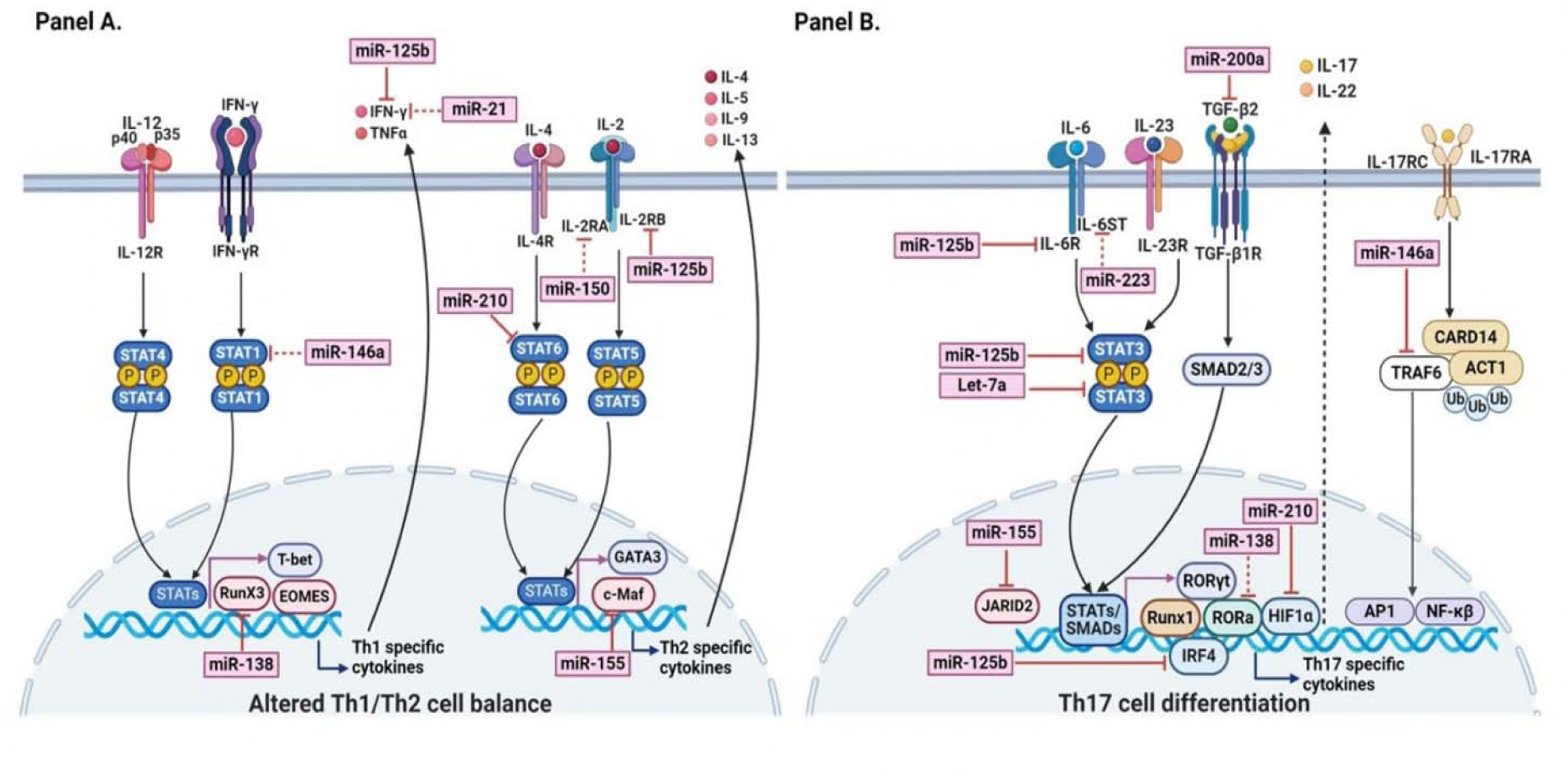
Panel. **A**. Working Model for miRNAs deciphered for their role in altered T cell subset differentiation. miRNAs are shown in rectangular boxes with validated targets represented with solid lines and predicted targets in dotted lines. 7/14 miRNAs with a role in psoriatic T cells are shown to impinge downstream targets that make a part of Th1/Th2 differentiation. **Panel B**. 8 /14 miRNAs are potentially involved in Th17 differentiation in psoriasis. (Figure created in BioRender. Madaan, P. (2024) https://BioRender.com/v63r892).

## 4. Discussion

A comprehensive search for T cell-specific miRNAs in the context of psoriatic T cells, gave a total of 14 miRNAs retrieved from 8 articles and 1 dataset of relevance, implying limited research findings available on miRNA-mediated dysregulation of T cell activation, survival, proliferation, and differentiation in psoriasis. Only 4/14 miRNAs viz miR-138, miR-200a, miR-210, and let-7a have been demonstrated to play a role in T cell proliferation and T cell subset differentiation based on the cognate target molecules and the altered T cell pathways [21, 23–26]. miR-138 targets the psoriasis-susceptibility gene Runt-related transcription factor 3 (RUNX3). Downregulation of miR-138 in CD4^+^ T cells leads to an increase in RUNX3 mediated Th1/Th2 ratio [21]. miR-200a expression level is shown to exhibit a statistically significant positive correlation with the Th17/Treg ratio along with the PASI score [25]. miR-210 with higher expression in CD4^+^ T cells from psoriasis vulgaris patients target FOXP3 and mediate Treg dysfunctionality along with promoting Th17 and Th1 cell differentiation over Th2 subset by inhibiting STAT6 and LYN [23, 24]. Th17-Th1 centric role of miR-210 is apparent from its therapeutic outcome in ameliorating imiquimod-induced psoriasis-like dermatitis in mice model [22]. Let-7a is another miRNA with lower expression in T cells from psoriatic patients with a role in T cell proliferation via targeting STAT3 [26].

The remaining 10/14 miRNAs viz miR-125b, miR-99a, miR-193b, miR-21, miR-17-92, miR-155, miR-146a, miR-150, miR-223, and miR-142-3p have been reported for their altered expression levels in T cells/T cell subsets with no direct proof of concept worked out to support their role in psoriasis as summarized in Table 1 [10, 27]. T cell preparation from psoriasis patients showed upregulation of miR-21, miR-125b, miR-99a, miR- 17 and miR-19 of miR-17-92 cluster and downregulation of miR-223 and miR-92 of miR-17-92 cluster [10, 38]. With respect to T cell subsets, miR-223 is documented to be upregulated in Th17 cells, while miR-21, miR- 146a, and miR-193b are shown to be downregulated [10]. In the Th1 subset, miR-99a and miR-193b are reported to be increased, while the Treg subset is demonstrated to have lower expression of miR-125b, miR-223, and miR-99a [10]. Interestingly, no change in expression of miR-155 and miR-142-3p is reported in T cells, Th17, Th1, and Treg despite their key role demonstrated in T cell activation and subset differentiation in health and other diseases [50, 60, 63, 85, 86]. Also, miR-150, shown to be downregulated in dermal infiltrates with no study on T cells/subsets, comprises one of the relevant miRNAs enlisted based on its involvement in the regulation of T cell development, activation, and CD4^+^ and CD8^+^ T cell subset differentiation [51–53]. Thus, expression analysis along with literature support for all the 10 miRNAs for their key role in maintaining T cell naivety, activation, proliferation, and/or T cell differentiation under homeostasis conditions prompted us to pick them as candidate miRNAs with potential implications in immune etiology of psoriasis. Interestingly upregulation of 5/10 miRNAs *i.e.,* miR-223, miR-142-3p, miR-155, miR-146a except miR-21 found in our study does not corroborate with earlier findings as sample type is different from the bulk CD3+ T cells used in our study along with the smaller sample size used in the previous studies. Nonetheless, the altered expression of these miRNAs signify their role in psoriasis.

Further, the network of integrated crosstalk of 15 signaling pathways derived, involved the validated and predicted targets downstream of the 14 miRNAs. Also gene ontology based molecular functions altered in psoriatic T cells comprised of signal transduction activating and inhibiting proteins. The findings on miRNA- mRNA target axes highlight their role in driving intricacies of T cell alterations in psoriasis. 3/15 pathways viz. T cell receptor signaling pathway, Th1/Th2 differentiation pathway, and Th17 differentiation pathway are well- studied T cell-specific pathways. Other pathways deciphered represent those involved in multiple cellular functionalities, including T cells.

T cell activation phenomenon is known to be associated with inflammatory landscape in psoriasis. An active T cell response with the psoriasis-specific secretion of pro-inflammatory cytokines like IFN-γ, TNF-α, and IL-17 is well documented in lesions as well as systemic circulation [87]. With limited studies on the activation status of T cells, lesional CD4^+^ and CD8^+^ active T cells are reported to be elevated in number with persistence of clonal T cell proliferation [88–90]. However, no change is reported for the transition of naïve to activated T cells in the peripheral circulation [91, 92]. In context of T cell activation associated pathways, T cell receptor (TCR) signaling is the key pathway for T cell effector functions involving multiple activating/inhibiting receptor-ligand interactions that fine-tune the downstream signaling cascades involving process-specific transcription factors like NFAT, AP-1, NF-κB with upstream adaptor proteins like LAT, GRb2, KRAS; pathway activating components comprising kinases like PI3K, AKT and inhibitory molecules for example phosphatases PTEN, SHIP1 [93]. Overall, TCR signaling functions as a key proximal signaling mechanism, subsequently leading to the activation of multiple distal signaling cascades, namely PI3K/AKT signaling, MAPK signaling, and mTOR signaling that together culminate in regulating gene expression involved in T cell activation/ survival/ proliferation and maintenance [94]. T-cell survival and proliferation readouts involve basic processes of cell cycle and apoptosis under homeostatic and disease conditions. The differential expression of cell cycle regulating transcription factors, different cyclins, and associated cyclin-dependent kinases (CDKs), along with molecules involved in extrinsic and intrinsic apoptosis pathways leading to caspase-mediated cell death, can determine T cell survival dynamics [95]. The p53 signaling comprises another pathway that mediates cell cycle arrest [96]. Together, these pathways integrate multiple signals to determine cell fate in terms of T-cell activation, survival, and proliferation.

Multiple miRNAs have been demonstrated to be differentially expressed and to regulate components of T cell receptor signaling, PI3K/MAPK/mTOR, cell cycle, apoptosis, and p53 signaling cascades. FOS, a component of AP-1 and NR2F6 that works as a repressor of NFAT and AP-1, has been shown to be targeted by miR-155 and miR-142-3p, respectively [49, 66, 97]. Negative regulators of T cell activation, CTLA-4 is targeted by miR- 155 and PD1 by miR-138, in contrast to PDK1, also targeted by miR-138, that promotes T cell activation [55, 56, 63]. Kinases like AKT3 and PIK3R1 involved in TCR-mediated signaling cascades like PI3K/AKT, mTOR, and MAPK are also validated targets of miR-150 and miR-21, respectively [53, 68]. Additionally, miR- 155 targets PIK3R1, a regulatory subunit of PI3K [65]. Other molecules with kinase activity, including mTOR, IGF1R, and IRAK1, the intermediate adaptor molecules like TRAF6, ARRB2 along with phosphatases like PTPN2, PTEN, SHIP1, and DUSP10 that regulate kinase activity, are all well documented as targets of different miRNAs as depicted in the result section, Table 2 and Table 3. TRAF6-IRAK1 components involved in the activation of MAPK pathways, such as p38 MAPK and JNK comprise direct targets of miR-146a [77, 98]. mTOR, the downstream target of miR-150 and miR-99a that comprises mTORC1 and mTORC2 complexes, enables T cells to transit from a quiescent state to an activated state by modulating their metabolism, survival, and migration [52, 99, 100].

RICTOR, a critical component of mTORC2, is also a direct target of miR-142-3p that modulates T-cell activation [48]. IGF1R targeted by miR-99a, also connects to PI3K/AKT and MAPK pathways [19]. ARRB2 is yet another molecule, a direct target of miR-150, which acts as a scaffold for various downstream TCR signaling molecules [54]. Specific phosphatases that regulate the phosphorylation events to fine-tune the T-cell activation also comprise targets of miRNAs of interest. PTPN2 and SHIP1 targeted by miR-155 have been shown to negatively regulate PI3K/AKT signaling-associated phosphorylation events in the context of CD8^+^ T cell activity [64]. On a similar note, miR-21, miR-223, and miR-17-92 are reported to regulate dephosphorylation events in the PI3K/AKT pathway by targeting PTEN and miR-21 targets MAPK phosphatase DUSP10 and regulate MAPK signaling [33, 71, 73, 80]. miR-193b-MYB, miR-21/miR-223-MEF2C, Let7a-Myc, and miR- 193b-KRAS represent other validated miR-target interactions that can affect T cell activation (Table 2 and Table 3) [59, 72, 74, 75, 101]. All miRNAs (13/14) except miR-210 can potentially modulate cell cycle, apoptosis, and related p53 pathway via multiple targets comprising cell cycle regulating transcription factors, different cyclins, and pro-apoptotic molecules (Table 2 and Table 3). E2F7 and E2F8, members of the E2F family of transcription factors, act as transcriptional repressors of cell cycle progression and have been validated as direct targets of miR-142-3p [50]. Cyclins act as crucial regulatory proteins that control cell cycle progression. Among these, CCND1, CCND3, and cyclin T1 (CCNT1) constitute validated targets with a significant role in G1/S transition. miR-193b and miR-138 targets CCND1 and CCND3 respectively, whereas CyclinT1 is a direct target of miR-150 and miR-223 [51, 57, 76]. Specific pro-apoptotic molecules have also been validated as targets of distinct miRNAs (Table 2 and Table 3). PDCD4, a direct target of miR-21 and miR-200a, along with Bim a direct target of miR-150, miR-155 and miR-17-92, work as pro-apoptotic factors [53, 67, 69, 79, 102].

Apart from all the experimentally validated targets downstream of the 14 miRNAs of interest in different cell types as per literature, a large number of predicted targets were also deciphered in our study based on the prediction tools with no experimental validation to date. These predicted targets majorly belong to signaling receptors viz ITPR3, GRa, ITGB8, FGFR2; transcription factors, e.g. NFAT5; intermediate signaling components such as GNB2, RASGRP1, NRAS, RAC1, RHEB; kinases, e.g. AKT1, RPS6KA1, PIK3CA and phosphatases like PPP2CA. In the context of cell cycle and apoptosis, other relevant signaling-associated molecules like cyclins, e.g., CCND2, anti-apoptotic proteins like BCL2, MCL-1, and pro-apoptotic molecules like CASP3, CASP7 comprise predicted targets of one or more miRNAs of interest. All the predicted/validated miRNA targets identified as components of T cell activation pathways with relevance in psoriasis needs to be further explored for their diagnostic and therapeutic value.

Importantly, miRNAs regulate T cell differentiation dynamics as well. T cell subset-specific quantitative and qualitative differences are well documented in health vs disease conditions. T cells comprise specific subsets with Th1, Th2, Th17, and Treg as the most explored with heterogeneous effector immune-functionalities required to counter diverse extrinsic and intrinsic challenges. Specific cytokine cues lead to the differentiation of naive T cells with molecular changes that define lineage commitment to different subsets involving differential expression of subset-specific transcription factors and effector cytokines [103]. Th1 cells are primarily involved in cell-mediated immunity, while Th2 cells modulate humoral immunity. IL-12 is a key cytokine that mediates the expression of T-bet as the master transcription factor responsible for Th1 differentiation [104]. In contrast, IL-4 induces GATA3, which serves as the master regulator of Th2 subsets [105]. Th17 cells are associated with inflammatory processes, while Tregs perform the balancing act with its anti-inflammatory role. IL-6, with a low dosage of TGF-β, leads to differential expression of lineage-defining master transcription factor RORγt inducing Th17-cell differentiation, whereas IL-2, in combination with TGF-β, induces Treg differentiation with FOXP3 as a key transcription regulator [103, 106]. In accordance with subset-specific cytokines and transcription factors involved, different signaling components together regulate their expression towards subset specific differentiation. Following the binding of different cytokines to their cognate receptors, the JAK/STAT pathway involving different STAT proteins mediates the expression of subset-specific transcription factors mentioned above. Multiple targets of miRNAs of interest, viz. lineage determining transcription factors, cytokine/cytokine receptors, chemokine/chemokine receptors, intermediate signaling activators, and inhibitors deciphered downstream of candidate miRNAs in our analysis, are related to T cell differentiation as clear from Table 2 and Table S2.

IL-12A/IL-12R and IFN-γ/IFN-γR cytokine/receptors mediate Th1 differentiating signaling pathways while IL- 4/IL4R with IL-2/IL-2R promote Th2 differentiation by activating subset-specific STAT proteins. STAT1 and STAT4 homodimerization leads to differential expression of Th1-specific transcription factor Tbet. Alternatively, STAT5 and STAT6 induce the expression of Th2 specific transcription factor GATA3 [107, 108]. RunX3 and c-Maf act as co-transcription factors in Th1 versus Th2 differentiation, respectively. miR-210 and miR-155 have been validated to target Th2 modulating STAT6 and c-Maf respectively and miR-138 is reported to target Th1 modulating RunX3 along with Th1 specific STAT1 as the predicted target of miR-146a [21, 23, 61]. The Wnt/β-catenin signaling pathway, an additional signaling mechanism in T cell subset differentiation, promotes Th2 differentiation over Th1 cells along with maintenance of CD8^+^ memory T cells [109]. The miR- targets deciphered in our study in association with Wnt signaling have not been experimentally validated. The key molecules, such as FZD4, APC, PRKACB, and ROCK2, RAC, constitute predicted targets downstream of miR-223 and miR-142-3p, representing components of Wnt signaling not yet explored. With respect to Th17 lineage commitment, IL-6/IL6R, together with TGF-β/TGFBR axes, trigger STAT-3 mediated Th17 specific expression of RORγt essential for Th17 lineage commitment [106, 110]. IRF4, HIF-1α, and RORα are other co-transcription factors facilitating Th17 differentiation along with RORγt [111].

Psoriasis is understood as a Th1/Th17-mediated disease with quantitative increase in Th1 and Th17 subsets in peripheral circulation as well as in skin lesions [6, 91, 112]. In line with this, Th17 signaling-specific validated targets were enriched downstream of candidate miRNAs viz. miR-125b targets STAT3 and IRF4 while miR-210 targets HIF-1α (Table 3) [58, 83, 84]. Jarid2, a DNA-binding protein, also represents one of the documented targets of miR-155 that recruits the Polycomb Repressive Complex 2 (PRC2) to chromatin and mediates repression of proinflammatory genes, thereby suppressing Th17 differentiation [62]. SOCS1, a direct target of miR-155, is another validated molecule retrieved that acts as a critical regulator of Th1 and Th17 differentiation [60]. RORA is an important predicted target downstream of miR-138 known to be involved in Th17 differentiation (Table 2). miR-125b can arrest T-cell differentiation by targeting IFNG, IL2RB, IL-10RA and IL6R [81, 82]. miR-200a is known to target TGFB2, albeit in the context of regulation of cell proliferation, migration, and invasion in renal cell carcinoma [78]. Along with these validated miRNA-targets enriched in our findings, multiple molecules that hold relevance in T cell subset differentiation viz IL-12A, IL-2RA, IL-2RB, IL-6ST, IL-17RD, and SMAD7 constitute predicted targets of miRNAs of interest (Table 2). Thus, candidate miRNAs deciphered can target subset-specific cytokine-receptor axes and modulate the dynamic process of T- cell differentiation in psoriasis.

In addition to cytokines, specific chemokine receptor signaling guide the migration of T cells to inflammatory sites, along with defining the functional characteristics of T cell subsets. CXCR3 and CCR5 define Th1; CCR3, CCR4, and CCR8 are Th2 specific and CCR6 is distinct to Th17 cells [113]. Among the miRNAs of interest, miR-21 is shown to target CCL20, which is a ligand to CCR6 and can modulate Th17 function, while CCL5 and CCL1 are retrieved as predicted targets of miR-146a and miR-21, respectively that are known to modulate Th1/ Th2 subsets [70]. All the validated miR-targets retrieved have been studied in different cellular contexts, viz T cell and non-T cell landscape. These, along with a large number of predicted targets, can potentially dysregulate T cell subset differentiation in psoriasis. Positioning of miRNAs with altered expression in psoriatic T cells in alignment with their known and predicted targets, unravel the possible molecular pathways that can drive altered T cell activation, proliferation, survival, and T cell differentiation. The findings provide a mechanistic base to validate identified miRNAs and their target molecule as potential diagnostic markers and precision therapeutics.

## 5. Conclusion

The study presents candidate miRNAs altered in psoriatic T cells with their possible downstream target mechanisms as the basis of psoriasis-specific T cell alterations. The identification of 14 differentially expressed miRNAs in psoriatic T cells suggests that miRNA dysregulation likely plays a significant role in T cell- mediated pathogenesis with widespread alterations in post-transcriptional gene regulation. However, the 14 miRNAs screened on the basis of their differential expression in psoriatic T cells need to be tested on a large patient cohort, since we retrieved limited literature and datasets that were relevant with respect to altered miRNAs in psoriatic T cells along with the small sample size used in the experimental work reported till date. Also, elucidation of 256 relevant targets as components of 15 T cell-associated pathways that can be dysregulated by the 14 miRNA candidates reveals a complex miRNA-mRNA interaction network potentially fine-tuned to give rise to an inflammatory T cell response in psoriasis. Our findings imply that multiple molecular modalities that regulate T cell functionality viz T cell receptor signaling, PI3K/AKT/ mTOR/MAPK cascades, cell cycle/survival pathways, and molecular checkpoints that regulate T cell subset differentiation can be modulated by specific miRNAs towards a psoriasis-specific T cell phenotype.

Scare studies on miRNA mediated altered T cell specific targets in psoriasis emphasizes a wide research gap in defining the real miRNA-target axes functional in dysregulated psoriatic T cells. However, the findings on miR-138, miR-210, and Let-7a and their validated targets in the context of psoriatic T cells confer high confidence on the miRNA-targets deciphered as potential mediators of T cell changes in psoriasis.

With this background, the proposed working models from our findings on how miRNAs may dysregulate T cell activation and differentiation offer testable hypotheses for further investigation underlining T cell-centric autoimmune changes in psoriasis. It is important to mention that rigorous proof of concept is required to support our findings since the miRNA-target effect is context-dependent and promiscuous in nature. Many of the targets retrieved in our *in silico* search are documented as validated miRNA targets in a setting of healthy individuals and/or other autoimmune diseases based on literature. Also, the downstream effector target for many of the select miRNAs are worked out in cell line models instead of primary T cell cultures from study subjects. Thus, the miRNA-target pathways projected in our findings provide a roadmap for future experimental studies that need to be carried out to access their role in disease etiology. Apart from better understanding the basis of T cell-driven disease manifestation, such studies on miRNA and their cognate targets can offer novel biomarkers and workable T cell centric therapeutic strategies in the field.

## Declarations

### Ethical approval and consent to participate

The study received formal ethical approval from AIIMS Bathinda (IEC/AIIMS/BTI/150), and the Central University of Punjab (CUPB/IEC/2016/045) with written informed consent obtained from all subjects.

### Data availability

The data generated or analyzed during this study are included in this published article (and its supplementary information files). Any further information regarding data generated in the current study can be availed from the corresponding author upon reasonable request.

### Author Contributions

MJ conceived the original idea. PM performed the in-silico analysis along with the literature review. NT, ST, and HRK helped with the miRNA-target network image. PM and MJ wrote the manuscript. DS and RT helped with the editing. MJ supervised and supported the research. All authors contributed to the article and approved the submitted version.

## Acknowledgement

Not Applicable

## Supplementary Information

The following supporting information is enclosed as supplementary. Table S1: List of GEO datasets explored for psoriasis-specific T cell-associated miRNAs; File S1: Venn Diagrams for 14 miRNAs showing the number of targets identified using four prediction tools miRDB, TargetScan, DianamicroT, and miRTarbase. Table S2: Additional targets for miR-17-92, other than those shown in Table 2, involved in specific pathways enlisted.

