## Supplementary File 1 for "Identification of T-Cell specific signature miRNAs and target pathways in Psoriasis"

**File S1:** Venn Diagrams for 14 miRNAs showing number of targets identified using four prediction tools miRDB, TargetScan, DianamicroT and miRTarbase.

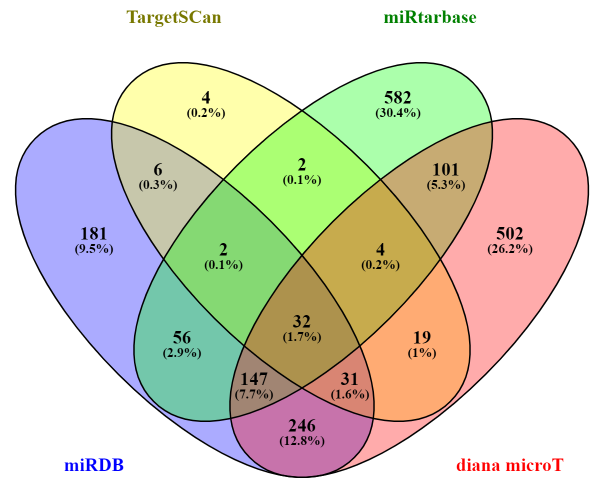

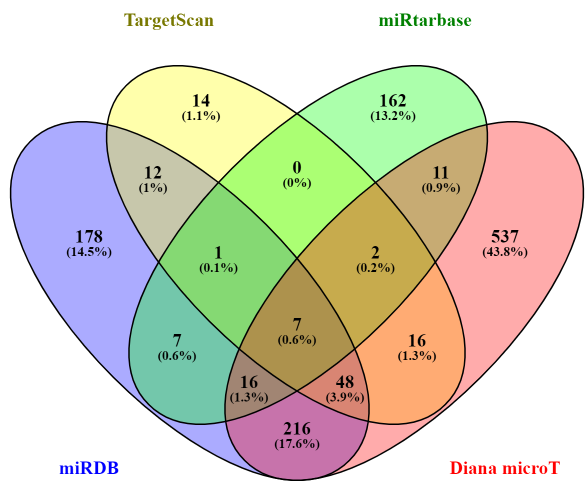

**miR-155	miR-146a**

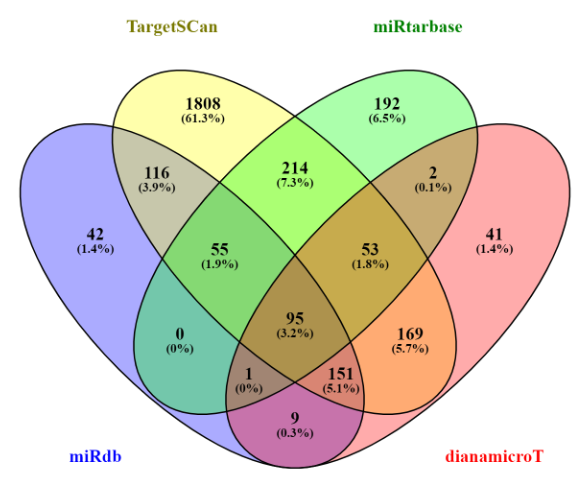

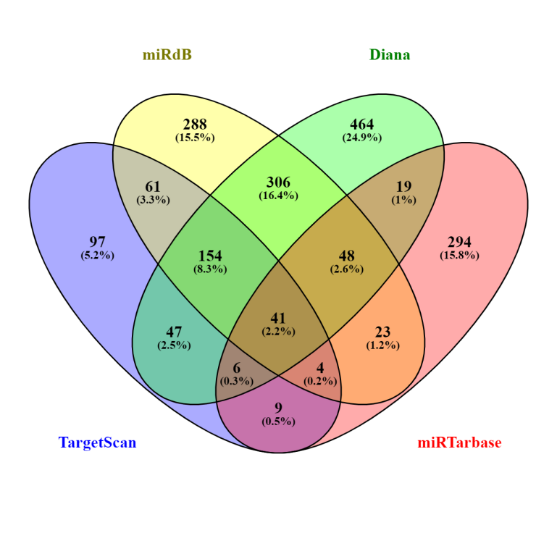

**miR-21**	**miR-125b-5p**

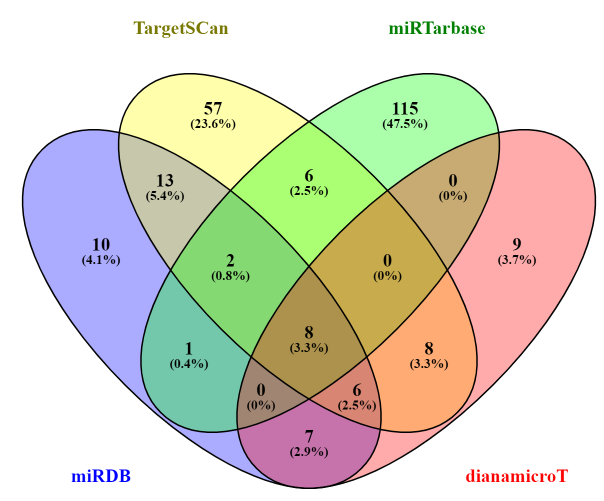

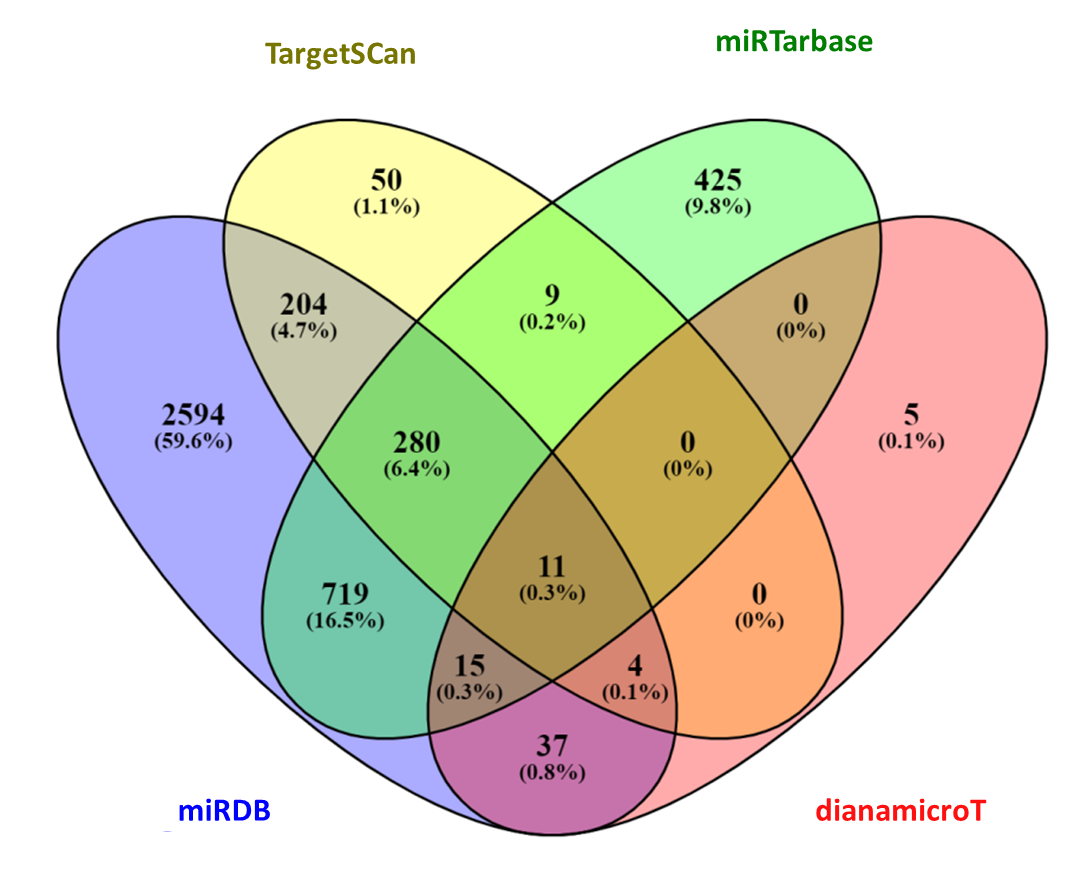

**miR-99a	miR-193b**

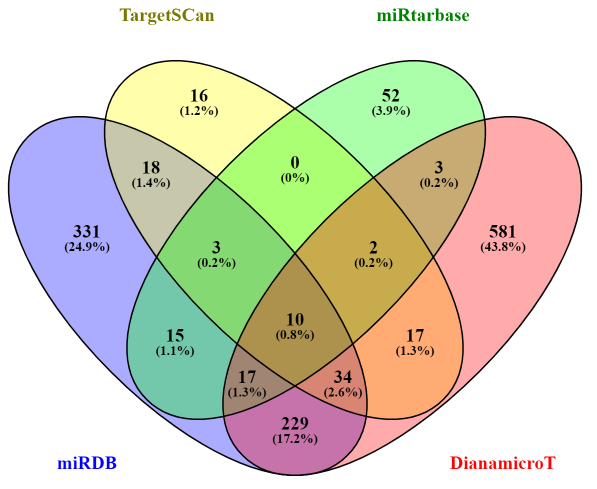

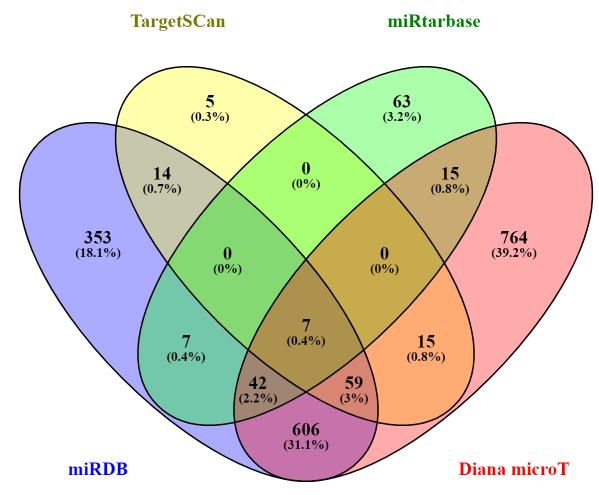

**miR-138a	miR-200a**

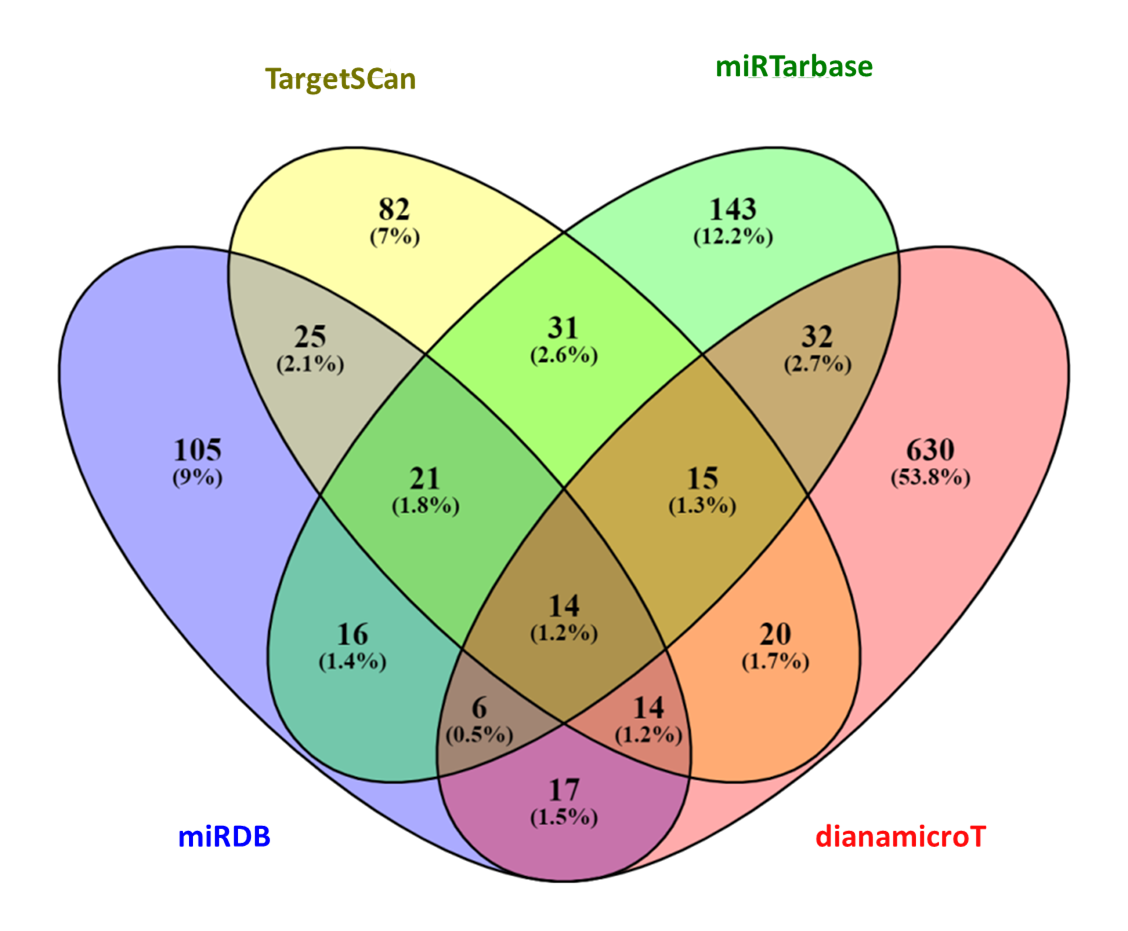

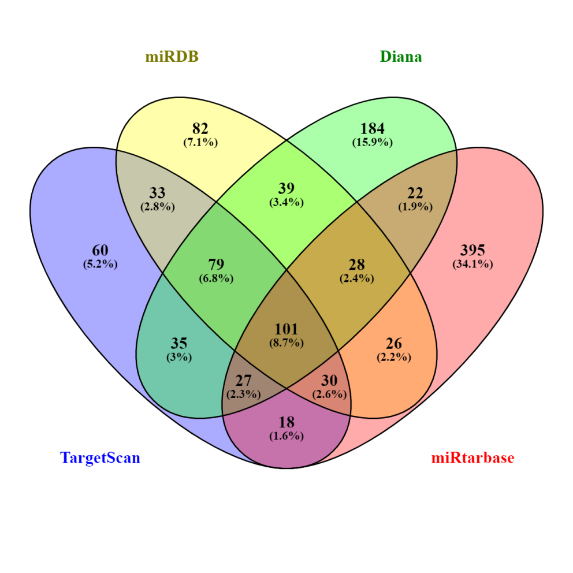

**miR-210	miR-142-3p**

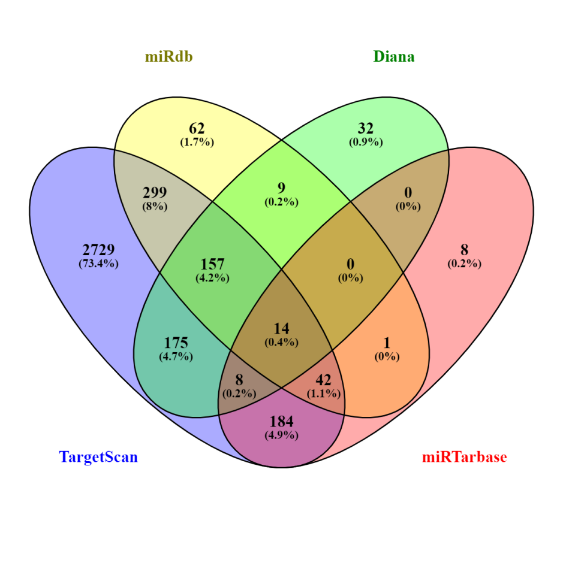

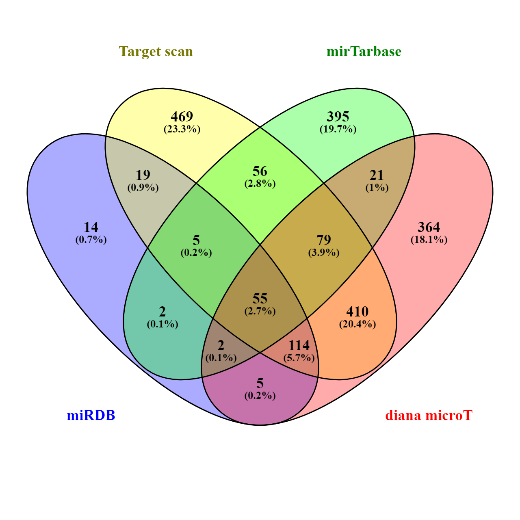

**miR-223	Let-7a**
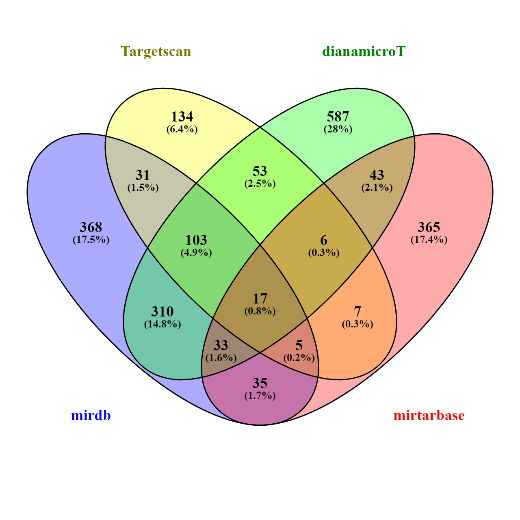

**miR-150**

**Venn diagrams showing common targets of members of miR-17-92 cluster**

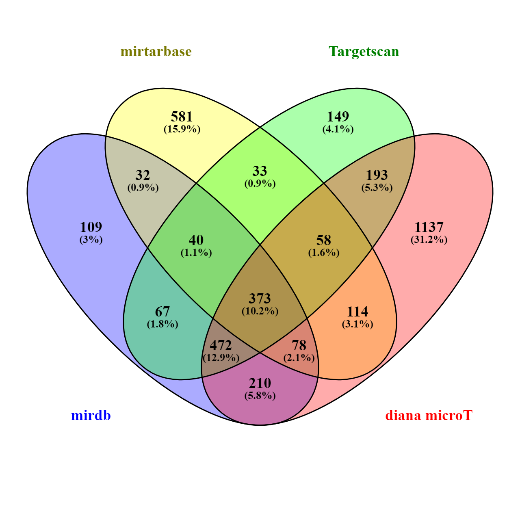

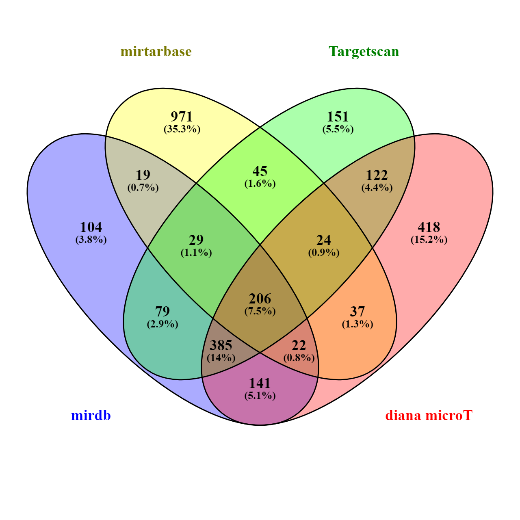

**miR-17-20a	miR-92a**

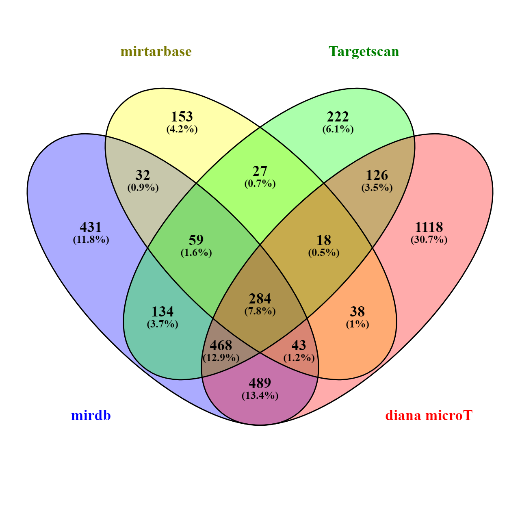

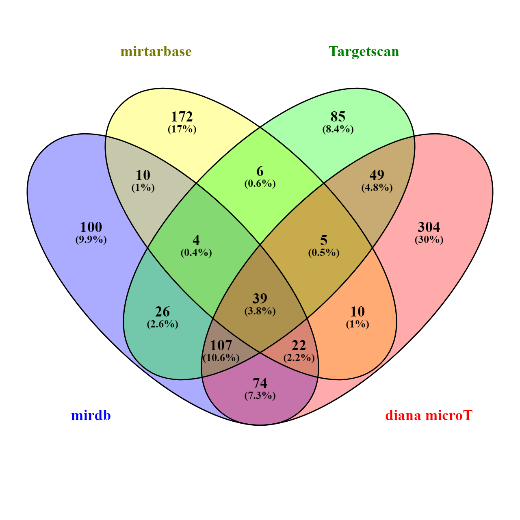

**miR-18a	miR-19a, 19b**
