## Supplementary Table 1 for "Identification of T-Cell specific signature miRNAs and target pathways in Psoriasis"

**Supplementary Table S1**

**Identification of T-Cell specific signature miRNAs and target pathways in Psoriasis**

**Priyanka Madaan ^1^, Nipanshi Tyagi ^2^, Sourabh Tyagi ^2^, Hemant Ritturaj Kushwaha ^2^ , Ravi Tandon ^2^ , Diya A’gitok Sangma^1^ and Manju Jain ^1^,***

1 Department of Biochemistry, Central University of Punjab, Village-Ghudda, Bathinda, Punjab, India;

^2^ School of Biotechnology, Jawaharlal Nehru University, New Delhi, India; (RT)

**Table S1:** List of GEO datasets explored for psoriasis-specific T cell-associated miRNAs

| **Sr. No** | **GEO Dataset** | **Link** | **Reference** |
| --- | --- | --- | --- |
| 1. | GSE 57012 | https://www.ncbi.nlm.nih.gov/geo/query/acc.cgi, GSE57012 | [3] |
| 2. | GSE 47598 | http://www.ncbi.nlm.nih.gov/geo/query/acc.cgi?acc=GSE47598 | [1] |
| 3. | GSE 103489 | https://www.ncbi.nlm.nih.gov/geo/query/acc.cgi, GSE103489 | [2] |
| 4. | GSE 55515 | https://www.ncbi.nlm.nih.gov/geo/query/acc.cgi, GSE55515 | [4] |
| 5. | GSE 78023 | https://www.ncbi.nlm.nih.gov/geo/query/acc.cgi?acc=GSE78023 | [5] |
| 6. | GSM871289 | https://www.ncbi.nlm.nih.gov/geo/query/acc.cgi?acc=GSM871289 | Unpublished data |
| 7. | GSE129373 | https://www.ncbi.nlm.nih.gov/geo/query/acc.cgi,GSE129373 | [6] |
| 8. | GSE142510 | https://www.ncbi.nlm.nih.gov/geo/query/acc.cgi?acc=GSE142510 | [7] |
