## Supplementary Table 2 for "Identification of T-Cell specific signature miRNAs and target pathways in Psoriasis"

**Supplementary Table S2**

**Identification of T-Cell specific signature miRNAs and target pathways in Psoriasis**

**Priyanka Madaan ^1^, Nipanshi Tyagi ^2^, Sourabh Tyagi ^2^, Hemant Ritturaj Kushwaha ^2^ , Ravi Tandon ^2^ , Diya A’gitok Sangma^1^ and Manju Jain ^1^,***

1 Department of Biochemistry, Central University of Punjab, Village-Ghudda, Bathinda, Punjab, India;

^2^ School of Biotechnology, Jawaharlal Nehru University, New Delhi, India; (RT)

| **miRNAs** | **miR-17-92 Targets** |
| --- | --- |
| **T cell receptor signaling** | RAF1, MAP3K8, MAPK14, PPP2R5E, PIK3R3, PPP3R1, MAPK1 |
| **PI3K/AKT signaling** | RAF1, BCL2L11, ERBB4, ITGA2, PPP2R5E, ITGA5, GNB5, FGF1, SGK3, CREB3L2, FASLG, MAPK1, THBS1, CDKN1A, RBL2, CHRM2, CCND1, IBSP, TSC1, ITGB8, PIK3AP1, SGK1, PIK3R3, JAK1, DDIT4, ERBB3, PRKAA1, CREB1, PHLPP2, STK11 |
| **mTOR signaling** | ULK1, RAF1, RRAGD, TSC1, SGK1, ATP6V1B2, CAB39, PIK3R3, GRB10, DDIT4, PRKAA1, WNT7B, STK11, MAPK1, CLIP1 |
| **MAPK signaling** | MAP2K3, TGFBR2, RAF1, MAP3K8, ERBB4, MAP3K1, MAP3K9, ELK4, MAP3K5, FGF1, MAPK1, MAPK14, HSPA8, MAP3K2, PRKACB, RAP1B, DUSP5, RAPGEF2, MAP2K4, RAP1A, ERBB3, PPP3R1, DUSP2, RPS6KA5, NF1 |
| **Apoptosis** | RAF1, BCL2L11, XIAP, MAP3K5, PIK3R3, DAB2IP, ATM, ITPR1, MAPK1 |
| **Cell cycle** | RB1, RAD21, RBL1, PPP2R5E, SMAD2, CDKN1A, PDS5B, RBL2, E2F5, ATM |
| **p53 signaling** | THBS1 RRM2 CDKN1A SESN3 ZMAT3 ATM |
| **Th1/Th2 differentiation** | MAPK14, JAK1, PPP3R1, MAPK1 |
| **Th17 differentiation** | TGFBR2, MAPK14, RARA, SMAD2, JAK1, RUNX1, PPP3R1, MAPK1 |
| **Wnt signaling** | BAMBI, PRICKLE2, DAAM2, MCC, PPP3R1, WNT7B |
| **JAK STAT signaling** | CDKN1A, RAF1, SOCS3, SOCS5, PIK3R3, SOCS6, JAK1 |
| **Cytokine cytokine interaction** | TGFBR2, TNFRSF2, TNFRSF12A, BMP2 |
| **IL-17 signaling** | MAPK14, MAPK1 |
| **TGF-β signaling** | THBS1, TGFBR2, BAMBI, RGMB, RBL1, ZFYVE9, TGIF, SMAD2, SMAD5, E2F5, BMP2, SKI, SKIL, MAPK1 |
| **Chemokine Receptor signaling** | RAF1, GNB5, PRKACB, RAP1B, PIK3R3, RAP1A, GRK6, MAPK1 |

**Table S2**: Additional targets for miR-17-92, other than those shown in Table 2, involved in specific pathways enlisted
